# Endothelial RRM2B-dependent mitochondrial DNA signalling drives doxorubicin-cardiotoxicity

**DOI:** 10.64898/2026.01.19.700350

**Authors:** Silvia Graziani, Altea Bonello, Raquel Brañas Casas, Giovanni Risato, Natascia Tiso, Francesco Argenton, Chiara Rampazzo, Satoko Shinjo, Margherita Zamberlan, Nicola Facchinello, Luca Scorrano, Giovanna Pontarin

**Affiliations:** Department of Biology, University of Padova, Padova, Italy; Department of Women’s and Children’s Health, University of Padova, Padova, Italy; Veneto Institute of Molecular Medicine, Padova, Italy; Department of Cell & Developmental Biology, Northwestern University Feinberg School of Medicine, Chicago, U.S.A; Department of Pharmacy and Biotechnology, University of Bologna, Bologna. Italy

## Abstract

Doxorubicin remains a mainstay of cancer therapy, yet its clinical use is constrained by severe cardiotoxicity, classically attributed to cardiomyocyte death. Here, we identify endothelial inflammation driven by mitochondrial DNA (mtDNA) turnover and release as a critical mediator of doxorubicin-induced cardiac injury. *In vivo*, a standard doxorubicin regimen activates the cGAS–STING innate immune pathway in endothelial cells before cardiomyocyte dysfunction or death occurs. Mechanistically, doxorubicin recruits the stress-inducible ribonucleotide reductase subunit RRM2B (p53R2) that normally sustains the mitochondrial deoxyribonucleotide pool, to promote aberrant mtDNA turnover and release, fuelling sterile inflammation. The FDA-approved BAX inhibitor Eltrombopag suppresses mtDNA release, attenuates endothelial inflammation, and prevents doxorubicin cardiotoxicity *in vivo* without compromising anticancer efficacy. These findings reveal an unrecognized proinflammatory role of RRM2B and establish endothelial inflammation as a central driver of chemotherapy-related cardiac injury, identifying the cardiac endothelium as a tractable target for cardioprotection during cancer treatment.

## Introduction

Doxorubicin (Dox) is a mainstay of therapy for haematological and solid malignancies in both adults and children. However, its potent antitumor efficacy is constrained by cumulative dose-dependent cardiotoxicity^1^. Cardiac injury can range from subclinical myocardial remodelling to overt cardiomyopathy, often emerging years after treatment and compromising long-term survival. Despite decades of research, the molecular and cellular mechanisms of Dox-induced cardiotoxicity (DIC) remain debated and effective cardioprotective strategies are lacking.

Mitochondrial toxicity is recognized as a key driver in DIC^2^. Dox accumulates in mitochondria through high-affinity binding to cardiolipin in the inner mitochondrial membrane, leading to dysfunction through excessive reactive oxygen species (ROS) generation, altered iron metabolism, and calcium imbalance. Dox also intercalates into mitochondrial DNA (mtDNA) in a dose dependent manner^3^, inducing supercoiling, torsional stress and nucleoid aggregation^4–7^. MtDNA instability is linked to Dox cardiotoxicity^8^: autopsy studies reveal reduced mtDNA content in human hearts, paralleling persistent mtDNA depletion and delayed-onset cardiomyopathy in Dox-treated mice^9–12^. Genetic models implicate mitochondrial topoisomerases in this process: cardiomyocyte-specific deletion of Top2β mitigates mitochondrial injury^13^, whereas Top1mt deficiency exacerbates mtDNA damage^14^. Moreover, Dox-mediated mtDNA activates the cyclic GMP–AMP synthase (cGAS)–stimulator of interferon genes (STING) innate immune pathway^7,15^, a signalling axis increasingly linked to cardiovascular disease and heart failure^16–18^.

While cardiomyocytes have been the traditional focus of investigation of the mechanism of Dox-toxicity, accumulating evidence identifies the vascular endothelium as an additional and likely underappreciated target. The cardiac endothelium preserves myocardial perfusion, regulates barrier integrity, modulates inflammation, and provides paracrine support to cardiomyocytes^19–21^. Endothelial toxicity, chronic vascular inflammation, and premature atherosclerosis have been reported in Dox-treated survivors^22^. In murine models, Dox reduces capillary density, increases microvascular permeability, and impairs angiogenesis ^23,24^. Notably, endothelial injury occurs largely independent of apoptosis, suggesting alternative mechanisms ^25–28^. Supporting this, endothelial-specific deletion of the autophagy effector Atg7 worsens ^29^, whereas loss of Sting ameliorates, Dox-induced cardiac dysfunction^30^, implicating mtDNA-driven sterile inflammation as a key contributor to endothelial dysfunction and DIC.

RRM2B, the stable small subunit of ribonucleotide reductase (RNR), provides deoxyribonucleotides for DNA repair and mtDNA maintenance in quiescent and differentiated cells^31^. It also supports DNA replication under hypoxia^32^ and scavenges ROS^33^, and is upregulated during cellular senescence^34^. Although constitutively expressed, RRM2B is transcriptionally induced by the tumor suppressor protein p53 in response to genotoxic stress such as Dox^35^. These properties suggest that RRM2B may modulate Dox-induced mitochondrial stress, mtDNA turnover, and sterile inflammation.

Here we identify endothelial RRM2B as a critical determinant of Dox cardiotoxicity. By sustaining mtDNA synthesis and activating the cGAS–STING pathway, RRM2B drives endothelial inflammation and cardiac injury. These findings uncover a previously unrecognized role for RRM2B and establish the cardiac endothelium as a tractable target for cardioprotection during cancer treatment.

## Results

### Transient exposure to Dox causes early vascular alterations in a zebrafish model

*Danio rerio* represents an effective model to study cancer therapy–associated cardiovascular toxicity, as it recapitulates cardiotoxic phenotypes and shares with humans conserved molecular signaling pathways^36^. We adapted a previously established zebrafish model for acute DIC^37^ by exposing embryos to various doses of Dox at 1 day post-fertilization (1 dpf), after cardiac development and the circulation onset. After 24 hours, the drug was removed, and embryos were maintained in drug-free water until 5 dpf for phenotyping (Extended Fig.1A).

Dox-exposed embryos developed normally but displayed transient, dose-dependent pericardial edema (Extended Fig. 1B-C). Cardiac contractility was irreversibly impaired, as shown by a dose-dependent reduction in heart rate (Fig. 1A, Extended Fig. 1D). Only the highest dose affected survival at the endpoint (Fig.1B, Extended Fig. 1E). Using the *Tg(tg:EGFP, myl7:EGFP)^ia300Tg^* transgenic line, which expresses GFP under the myosin light chain promoter enabling high-speed cardiac imaging, we found that even the lowest dose of Dox increased atrial diastolic area and reduced ventricular fractional shortening (Fig.1C-E). Microscopy of the *Tg(kdrl:GFP)^s843Tg^* line, which labels endothelial cells, revealed that Dox narrowed the dorsal aorta (DA) and thinned the intersegmental vessels (ISVs) compared with control (DMSO-treated) individuals, indicating vascular alterations after Dox exposure (Fig.1F-G, Extended Fig. 1F-G). Notably, these vascular defects preceded detectable cardiac dysfunction.

**Figure 1.**
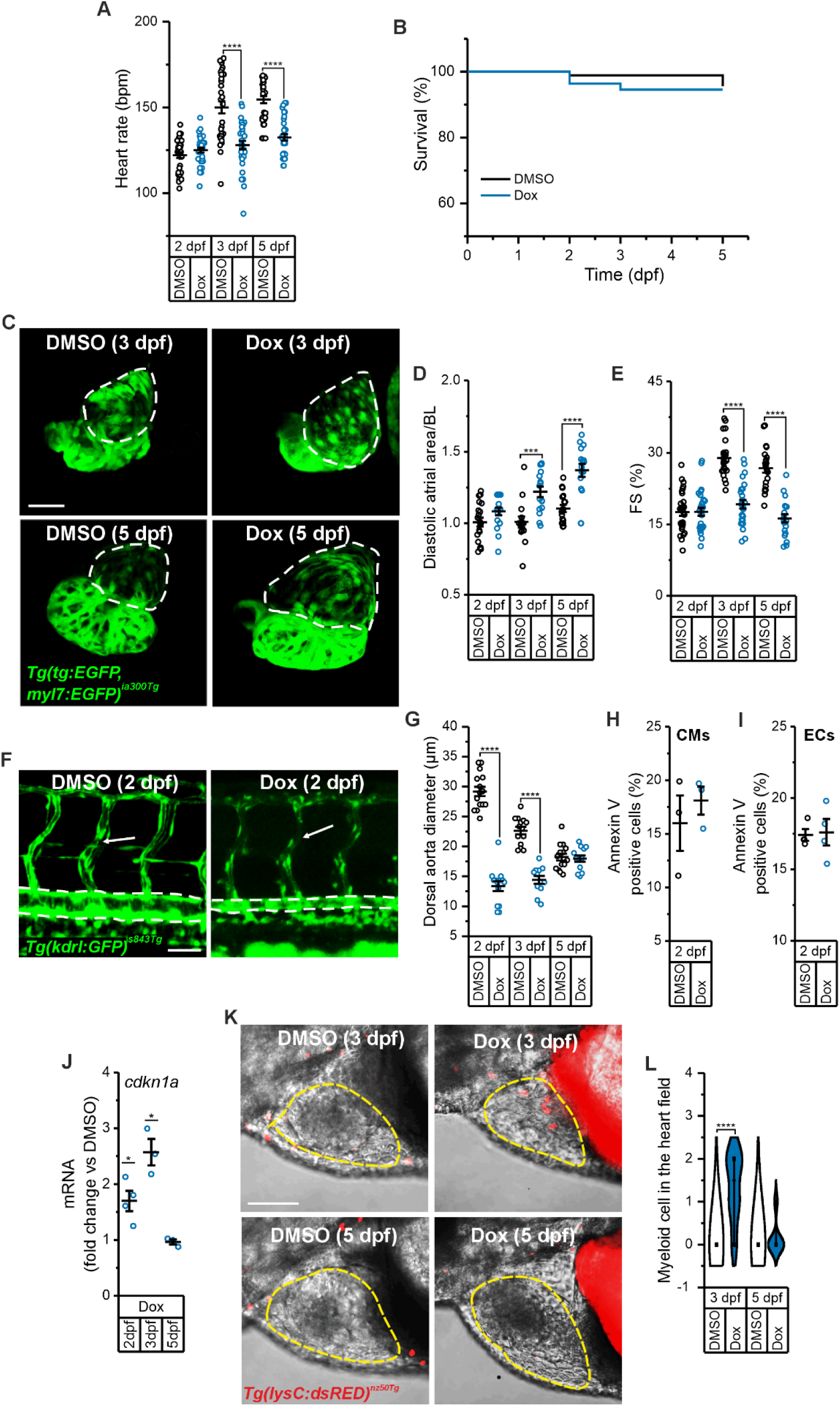
Zebrafish model faithfully recapitulates doxorubicin-induced cardiotoxicity. **A.** Heart rate analysis at different dpf in Dox-treated embryos and DMSO controls. Bpm, beats per minute. DMSO (n=34), 10 μM Dox (n=35) at 2 dpf, DMSO (n=33), 10 μM Dox (n=32) at 3 dpf, DMSO (n=35), 10 μM Dox (n=34) at 5 dpf. N=3 independent experiments. **** p<0.0001 in a two-sample t-test with Welch’s post hoc analysis for mean comparison or in a Mann-Whitney test. **B.** Kaplan-Meier Survival analysis curves of Dox-treated embryos and DMSO controls. DMSO (n=29), 10 μM Dox (n=30) at 2 dpf, DMSO (n=30), 10 μM Dox (n=27) at 3 dpf, DMSO (n=29), 10 μM Dox (n=24) at 5 dpf. N=3 independent experiments. p ≥ 0.05 in a log-rank test. **C.** Representative confocal images of atrial diastolic phase in Dox-treated and control 3 and 5 dpf *Tg(tg:EGFP, myl7:EGFP)^ia300Tg^* embryos. Dotted lines contour the atrium. Scale bar 50 µm. **D.** Quantification of diastolic atrial area in experiments as in C. DMSO (n=20), 10 μM Dox (n=18), at 2 dpf, DMSO (n=15), 10 μM Dox (n=16) at 3 dpf, DMSO (n=18), 10 μM Dox (n=16) at 5 dpf. N=3 independent experiments. *** p<0.001; **** p<0.0001 in a two-sample t-test with Welch’s post hoc analysis for mean comparison**. E.** Quantification of fractional shortening (FS) in experiments as in C. DMSO (n=29), 10 μM Dox (n=28) at 2 dpf, DMSO (n=26), 10 μM Dox (n=26) at 3 dpf, DMSO (n=21), 10 μM Dox (n=20) at 5 dpf. N=3 independent experiments. **** p<0.0001 in a two-sample t-test with Welch’s post hoc analysis for mean comparison. **F.** Representative confocal images of the trunk vasculature in Dox-treated and control 2 dpf *Tg(kdrl:GFP)^s843Tg^* embryos. Arrows indicate the intersegmental vessels (ISVs), dotted lines contour the dorsal aorta (DA). Scale bar 50 µm. **G.** Quantification of DA diameter in experiments as in F. DMSO (n=15), 10 μM Dox (n=14) at 2 dpf, DMSO (n=14), 10 μM Dox (n=12) at 3 dpf, DMSO (n=14), 10 μM Dox (n=13) at 5 dpf. N=3 independent experiments. **** p<0.0001 in a two-sample t-test with Welch’s post hoc analysis for mean comparison or in a Mann-Whitney test. **H-I.** Average ± SEM of Annexin-V+, GFP+ cardiomyocytes, CMs (H), and endothelial cells, ECs (I), isolated from 2 dpf *Tg(tg:EGFP, myl7:EGFP)^ia300Tg^* and *Tg(kdrl:GFP)^s843Tg^* embryos treated with DMSO or 10 μM Dox. N=3-4 independent experiments. p ≥ 0.05 in a two-sample t-test with Welch’s post hoc analysis for mean comparison. **J.** Expression level of *cdkn1a* at different dpf determined by qRT-PCR in GFP+ ECs sorted from *Tg(kdrl:GFP)^s84Tg3^* embryos treated as indicated. N=3-4 independent experiments. * p<0.05 in a two-sample t-test with Welch’s post hoc analysis for mean comparison. **K.** Representative confocal image of the heart and myeloid cells in embryos from the transgenic line *Tg(lysC:dsRed2)^nz50Tg^* at 3 and 5 dpf. Red dots indicate the dsRED+ myeloid cells, while dotted lines contour the heart. Scale bar 100 µm. **L.** Quantification of myeloid cells in fish heart. DMSO (n=18), 10 μM Dox (n=18) at 3 dpf, DMSO (n=18), 10 μM Dox (n=18) at 5 dpf. N=4-5 independent experiments. **** p<0.0001 in a One-sample Wilcoxon signed rank test.

To assess whether Dox-induced cardiovascular defects were linked to cell death we performed Annexin-V staining on sorted GFP-positive cardiomyocytes (CMs) and endothelial cells (ECs). Apoptosis occurred only at the highest Dox dose (Fig.1H-I, Extended Fig.1H-I).

Quantitative PCR (qRT-PCR) analysis revealed induction of the cell cycle regulator Cyclin Dependent Kinase Inhibitor 1A (*cdkn1a*) in ECs, indicating non-lethal endothelial responses alongside cardiomyocyte death (Fig.1J).

Together, these findings show that transient Dox exposure induces early endothelial injury and sustained myocardial dysfunction, even in the absence of overt cell death, with vascular alterations preceding cardiomyocyte damage.

### Dox induces cGAS–STING activation in endothelial cells

Inflammation is a hallmark of DIC. In the *Tg(lysC:dsRED)^nz50Tg^* line, which marks myeloid cells, Dox exposure led to transient macrophage and neutrophil accumulation within the myocardium, consistent with cardiac inflammation (Fig.1K-L). The endothelium regulates immune cell trafficking and acts as a gatekeeper of tissue inflammatory responses. Aberrant activation of different DNA-sensing pathways by Dox, including the NOD-, LRR- and pyrin domain-containing protein 3 (NLRP3) inflammasome, Toll-like receptor 4 (TLR4), the cGAS)–STING) pathway, and Neutrophil Extracellular Trap (NET) formation^38^, has been primarily investigated in cardiomyocytes and myeloid cells, while endothelial contributions to sterile inflammation remain largely unexplored.

In our *in vivo* model of DIC, mRNA profiling of GFP-sorted ECs at different timepoints revealed that Dox triggered a transient, broad inflammatory response, with early induction of Interleukin 1 beta (*il1b)*, Interleukin 6 (*il6*) and Interferon Regulatory Factor 3 (*irf3*) followed by interferon phi 3 (*ifnphi3*), the zebrafish orthologue of interferon alpha 1, 10 and 13 (Fig. 2A). These results are consistent with activation of inflammasome and cGAS-STING signalling. Dox also induced *ifnphi3* and *irf3* in GFP-sorted cardiomyocytes, but not *il6* and *il1b*, indicating that the immune signaling profiled here reflected a cell type-specific response (Extended Fig. 2A). To dissect the relative contribution of different innate immune pathways in DIC, we cotreated fish with Dox and the STING inhibitor H151. H151 prevented cardiac myeloid cells accumulation and preserved heart function, as indicated by normal heart rate and atrial diastolic area (Fig. 2B-E). H151 selectively inhibited *irf3* and *ifnphi3*, but not *il6* or *il1b* induction, demonstrating that cGAS–STING activation in Dox-treated ECs is uncoupled from NF-κB (Nuclear factor kappa-light-chain-enhancer of activated B cells) dependent transcription, resembling chronic mtDNA stress responses (Fig. 2A).

**Figure 2.**
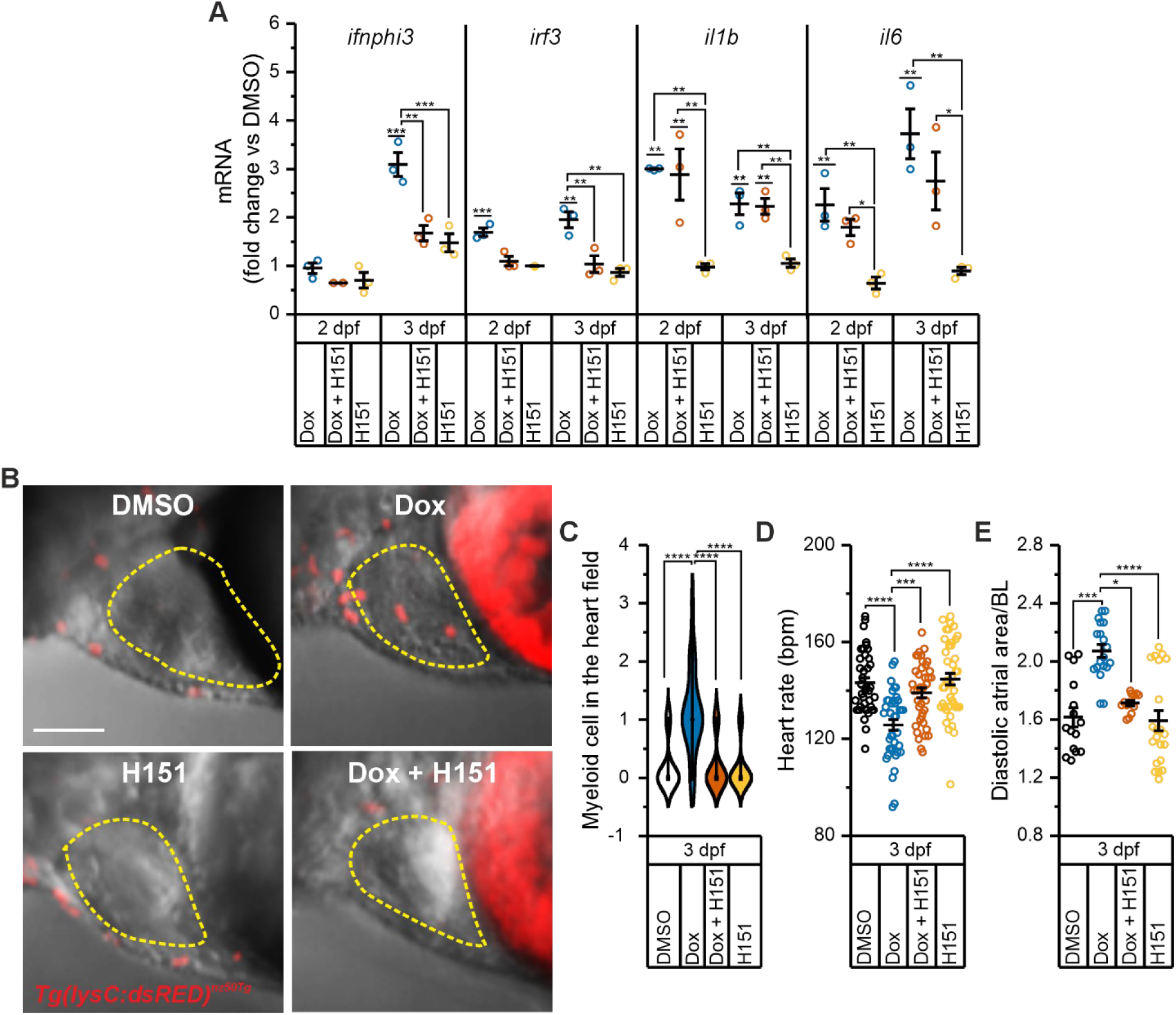
Inhibition of STING protects from Doxorubicin-induced cardiotoxicity *in vivo*. **A.** qRT-PCR time course analysis of the indicated genes in GFP+ ECs sorted from *Tg(kdrl:GFP)^s843Tg^* embryos treated as indicated. N=3 independent experiments. * p<0.05; ** p<0.01; *** p<0.001 in a one-way ANOVA test with Tukey’s post hoc analysis for mean comparison. **B.** Representative confocal image of the heart and myeloid cells in embryos from the transgenic line *Tg(lysC:dsRed2)^nz50Tg^* at 3 dpf. Red dots indicate the dsRED+ myeloid cells, while dotted lines contour the heart. Scale bar 100 µm. **C.** Quantification of myeloid cells in fish heart. DMSO (n=22), Dox (n=21), Dox + H151 (n=22), H151 (n=22). N=3 independent experiments. **** p<0.0001 in a Kruskal-Wallis ANOVA test with Dunn’s post hoc analysis for mean comparison. **D.** Heart rate analysis in embryos treated as indicated at 3 dpf. Bpm, beats per minute. DMSO (n=41), 10 μM Dox (n=39), 10 μM Dox + 2 μM H151 (n=39), 2 μM H151 (n=42). N=3 independent experiments. *** p<0.001; **** p<0.0001 in a one-way ANOVA test with Tukey’s post hoc analysis for mean comparison. **E.** Quantification of diastolic atrial area in the transgenic line *Tg(tg:EGFP, myl7:EGFP)^ia300Tg^* treated as indicated at 3 dpf. DMSO (n=15), 10 μM Dox (n=19), 10 μM Dox + 2 μM H151 (n=13), 2 μM H151 (n=21). N=3 independent experiments. * p<0.05; *** p<0.001; **** p<0.0001 in a Kruskal-Wallis ANOVA test with Dunn’s post hoc analysis for mean comparison.

Collectively, these results show that Dox induces cardiac inflammation associated with cGAS–STING activation in both endothelial cells and cardiomyocytes, and that STING inhibition protects against Dox-induced cardiac dysfunction.

### Dox-induced RRM2B activity drives mitochondrial DNA expansion in human endothelial cells

To elucidate the molecular mechanism driving cGAS-STING activation, we exposed human cardiac microvascular endothelial cells (HCMECs) to a clinically relevant sublethal dose of Dox and analysed them after drug withdrawal (Extended Fig. 3A). Similar to HUVECs^28^, Dox treatment induced cell cycle arrest and senescence, by Cyclin A loss, sustained p21^Waf/Cip1^, reduced Lamin B1 level and accumulation of senescence-associated β-galactosidase (SA-β-gal) positive cells (Extended Fig. 3B-C).

Because Mitochondrial DNA (mtDNA) is a Dox target and a potent cGAS-STING agonist^39^, we examined mtDNA dynamics in HCMECs and HUVECs after treatment. qRT-PCR revealed a progressive increase in mtDNA copy number per cell following Dox exposure (Fig. 3A, Extended Fig. 3D). BrdU pulse-chase assays in HUVECs 8 days after Dox exposure revealed that mtDNA synthesis exceeded degradation, indicating an imbalance favouring mtDNA accumulation (Fig. 3B-C). We next examined pathways supplying mitochondrial deoxynucleoside triphosphates (dNTPs). In non-proliferating cells, dNTPs are generated through the cytosolic *de novo* pathway mediated by p53-inducible RRM2B and the mitochondrial salvage pathway involving thymidine kinase 2 (TK2) and deoxyguanosine kinase (dGUOK). After Dox exposure, *TK2* remained unchanged, *dGUOK* decreased and RRM2B expression increased in a p53-dependent manner, while canonical RRM2 was lost (Fig. 3D-F, Extended Fig. 3E-G). Inhibition of RNR with hydroxyurea (HU) after RRM2 loss (i.e., 1 day after Dox removal) abrogated mtDNA expansion in both endothelial cell types (Fig. 3A, Extended Fig. 3D), underscoring the essential role of RRM2B-driven *de novo* synthesis in sustaining mtDNA accumulation.

**Figure 3.**
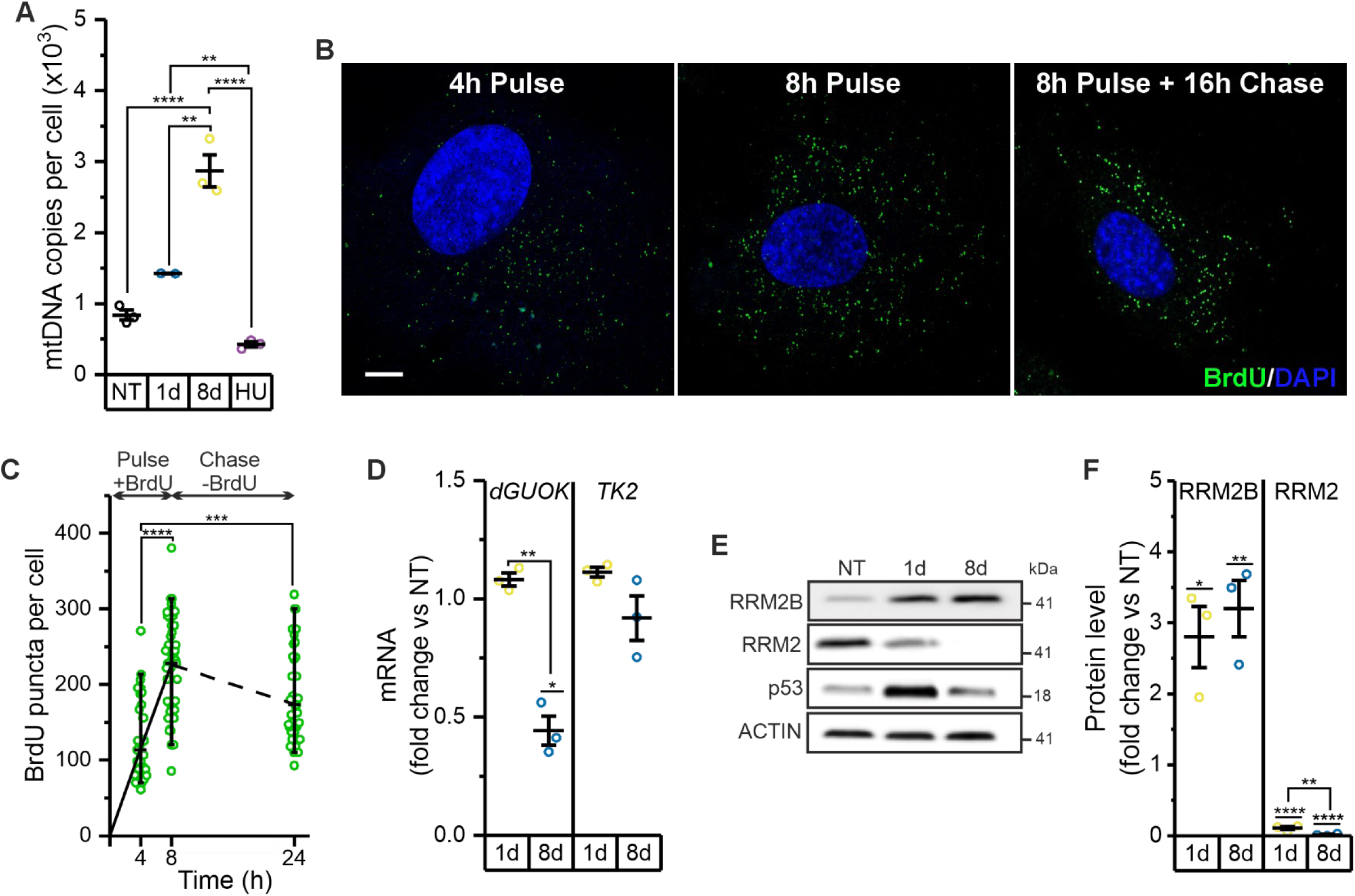
Doxorubicin induces RRM2B to sustain mitochondrial DNA expansion in endothelial cells. **A.** Quantification of mtDNA copy per cell determined by qPCR in HUVECs, not treated (NT), during Dox exposure (1d) and after drug withdrawal (8d). Hydroxyurea (HU) was added one day after Dox withdrawal. N=3 independent experiments. ** p<0.01; **** p<0.0001 in a one-way ANOVA test with Tukey’s post hoc analysis for mean comparison. **B-C.** Time curve of Bromodeoxyuridine (BrdU) incorporation into mtDNA in HUVECs after drug withdrawal (8d). HUVECs were pulsed with 10 µM BrdU (4h-8h) and chase in BrdU-free medium (16h), followed by immunostaining with anti-BrdU antibody (green puncta) to visualize newly synthesized mtDNA and counterstained with DAPI (blue signal) to detect cell nucleus. Representative images (B) and quantification (C) of number of green puncta/cell for each pulse (continuous line) and chase (dotted line) time. Scale bar 10 µm. N=3 independent experiments. *** p<0.001; **** p<0.0001 in a Kruskal-Wallis ANOVA test with Dunn’s post hoc analysis for mean comparison. **D.** Expression level of *dGUOK* and *TK2* by qRT-PCR in HUVECs, during Dox exposure (1d) and after drug withdrawal (8d). TK2, thymidine kinase 2. dGUOK, deoxyguanosine kinase. N=3 independent experiments. * p<0.05; ** p<0.01 in a one-way ANOVA test with Tukey’s post hoc analysis for mean comparison. **E.** Representative immunoblot image and **F.** quantification of the canonical RRM2 and alternative p53 inducible RRM2B small RNR subunit in HUVECs, not treated (NT), during Dox exposure (1d) and after drug withdrawal (8d). N=3 independent experiments. * p<0.05; ** p<0.01; **** p<0.0001 in in a one-way ANOVA test with Tukey’s post hoc analysis for mean comparison.

Quasi super-resolution Airyscan imaging of Dox-treated ECs revealed dsDNA positive dots in the cytosol, not colocalizing with mitochondria that appeared elongated and highly interconnected. Notably, we could not detect any cytosolic chromatin fragments (CCFs) in Dox-induced senescent ECs^40^ (Fig. 4A–C; Extended Fig. 3H–J). qPCR and immunoblot analyses confirmed the mtDNA nature of the increased cytosolic DNA and enrichment of the core mtDNA nucleoids components Transcription Factor A, Mitochondrial (TFAM), indicating mitochondrial DNA leakage (Fig. 4D–F). Thus, endothelial mitochondria chronically release mtDNA into the cytosol after the first Dox insult.

**Figure 4.**
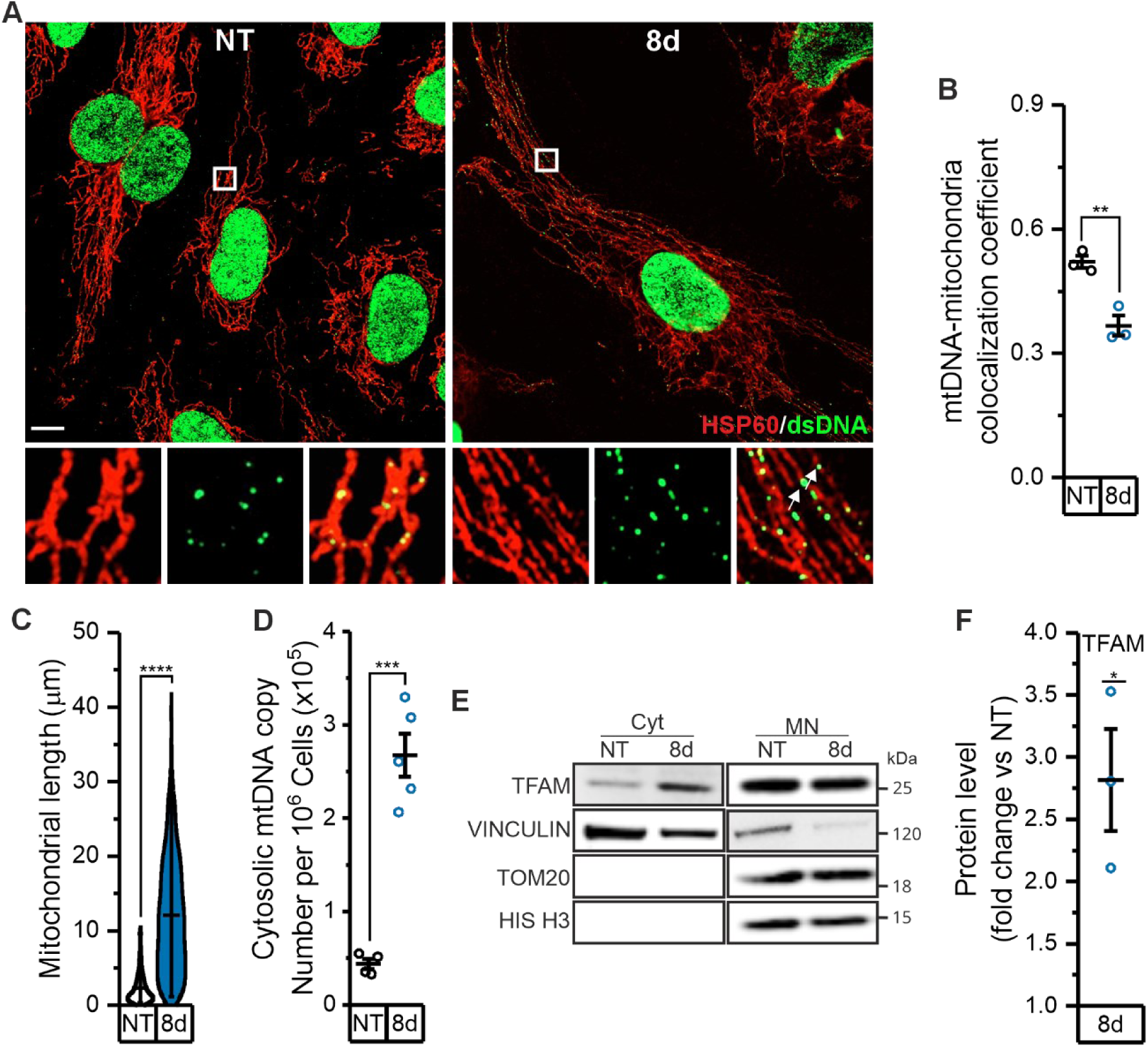
Doxorubicin-treated endothelial cells chronically release mtDNA in the cytosol. A-C. Immunostaining of HUVECs not treated (NT) and after Dox withdrawal (8d). **A.** Representative super-resolution Airyscan confocal images of dsDNA (green) and HSP60 (red). Scale bar 10 µm. At bottom, the magnified images show mtDNA (green) and mitochondria (red), showing that most DNA foci are located within mitochondria with some foci in the cytoplasm of Dox treated HUVECs. Arrows indicate cytosolic dsDNA signals. **B.** Quantification of the colocalization coefficient between mtDNA and mitochondria. N=3 independent experiments. ** p<0.01 in a two-sample t-test with Welch’s post hoc analysis for mean comparison. **C.** Quantification of mitochondrial length. NT (n=963), 8d (n=1027). N=3 independent experiments. **** p<0.0001 in in a Mann-Whitney test. **D.** mtDNA copy number per million cells determined by qPCR analysis in cytosolic fraction isolated from HUVECs not treated (NT) and after Dox withdrawal (8d). N=4/5 independent experiments. *** p<0.001 in a two-sample t-test with Welch’s post hoc analysis for mean comparison. **E.** Representative immunoblot image and **F.** quantification of TFAM present in the cytosolic (Cyt) and mitochondrial/nuclear (MN) fraction isolated from HUVECs not treated (NT) and after Dox withdrawal (8d). Vinculin, TOM20 and HIS H3 were used to assess the absence of cytosolic, mitochondrial and nuclear cross-contamination among the different fractions. N=3 independent experiments. * p<0.05 in one-sample t-test.

### RRM2B activity sustains cGAS-STING-inflammatory response

We next assessed whether mtDNA expansion and release drive inflammatory gene expression. Endothelial cells exhibited a biphasic cytokine response, with early Interleukin-1 beta (*IL-1B*) induction and later C-X-C motif chemokine ligand 10 (*CXCL10*) and Interleukin-8 (*IL-8*) expression (Fig. 5A). Depletion of mtDNA (ρ⁰ cells) abolished chronic cytokine induction without affecting senescence (Fig. 5B), showing that mtDNA is required for sustained inflammation. Inhibition of RNR with HU suppressed *CXCL10* but not *IL-8* induction, indicating that RRM2B activity selectively sustains cGAS-STING dependent signalling (Fig. 5C; Extended Fig. 3K). Treatment with the STING inhibitor H151 produced identical results, confirming that RRM2B promotes mtDNA-dependent, but NF-κB independent, inflammation. This profile mirrors that induced by chronic cytosolic mtDNA release in Tfam^+/-^ MEFs^15^ and by Dox in iPSC-derived or neonatal cardiomyocytes^7^.

**Figure 5.**
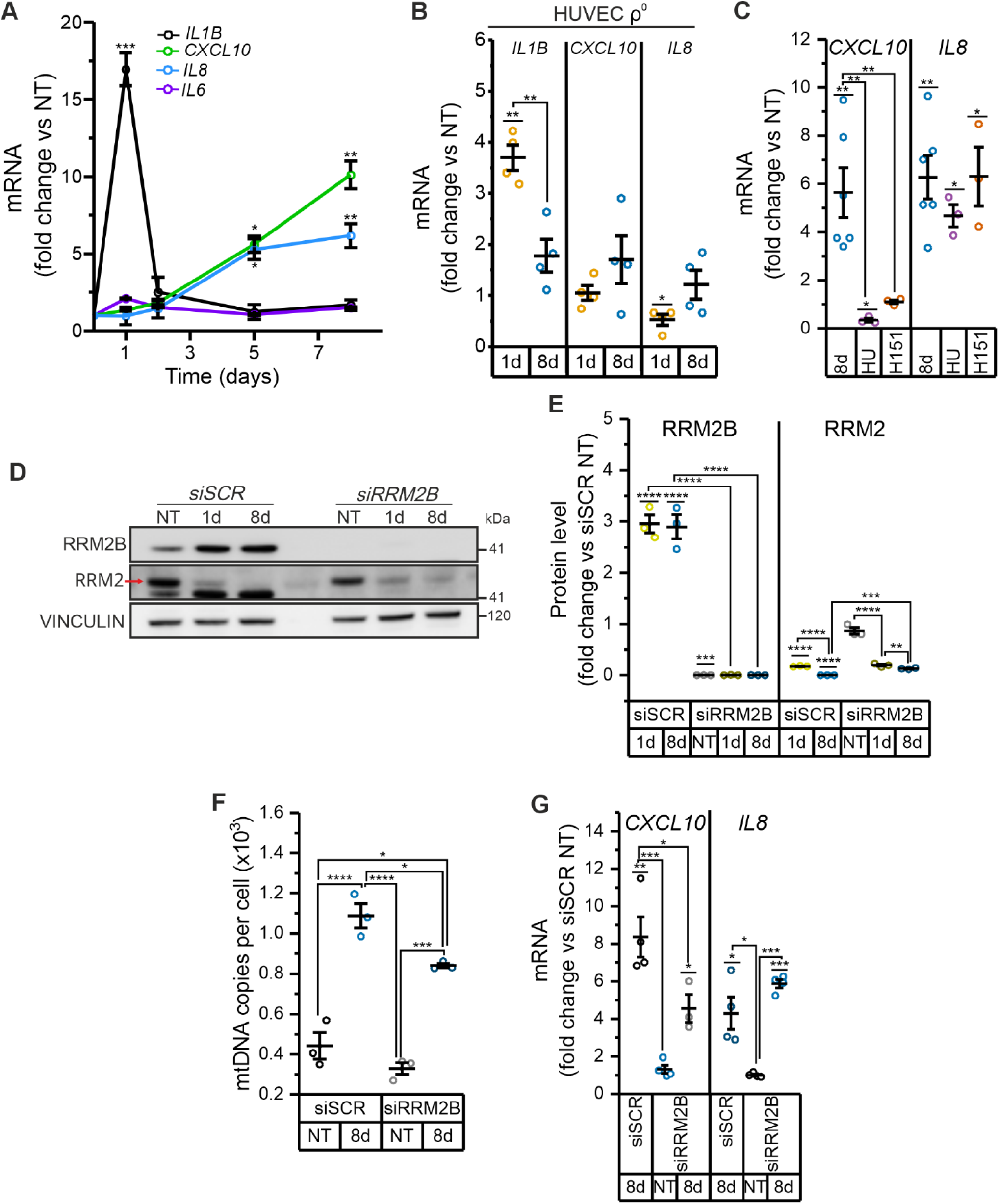
RRM2B couples doxorubicin-induced mitochondrial stress to cGAS–STING activation in endothelial cells. **A.** qRT-PCR time course analysis of the indicated inflammatory genes in HUVECs not treated (NT), during Dox exposure (1d) and after drug withdrawal (2, 5, 8d). N=3 independent experiments. * p<0.05; ** p<0.01; *** p<0.001 in a one-way ANOVA test with Tukey’s post hoc analysis for mean comparison. **B.** qRT-PCR time course analysis of the indicated inflammatory genes in mtDNA-depleted HUVECs (HUVECs ρ^0^) not treated (NT), during Dox exposure (1d) and after drug withdrawal (8d). N=4 independent experiments. * p<0.05; ** p<0.01 in a one-way ANOVA test with Tukey’s post hoc analysis for mean comparison. **C.** Expression levels of indicated genes measured by qRT-PCR in HUVECs, not treated (NT), and after drug withdrawal (8d). Hydroxyurea (HU) or Sting inhibitor H151 (H151) was added one day after Dox withdrawal. N=3/6 independent experiments. * p<0.05; ** p<0.01 in a one-way ANOVA test with Tukey’s post hoc analysis for mean comparison. **D-G.** RRM2B silencing in HUVECs. HUVECs were transfected with either a negative control siRNA (siSCR) or RRM2B targeting siRNAs (siRRM2B). Analysis was performed in cells not treated (NT), during Dox exposure (1d) and after drug withdrawal (8d). **D.** Representative immunoblot blot images and **E.** quantification of RNR subunit RRM2B and RRM2. N=3 independent experiments. ** p<0.01; *** p<0.001; **** p<0.0001 in a one-way ANOVA test with Tukey’s post hoc analysis for mean comparison. **F.** mtDNA copy number per million cells determined by qPCR analysis. N=3 independent experiments. * p<0.05; *** p<0.001; **** p<0.0001 in a one-way ANOVA test with Tukey’s post hoc analysis for mean comparison. **G.** Expression levels of indicated genes measured by qRT-PCR. N=3/4 independent experiments. * p<0.05; ** p<0.01; *** p<0.001 in a one-way ANOVA test with Tukey’s post hoc analysis for mean comparison.

RRM2B silencing (siRRM2B) in HUVECs reduced both mtDNA copy number and *CXCL10* induction, while *IL-8* remained unaffected (Fig. 5D-G). Given that RRM2 levels are substantially higher than RRM2B in cycling cells^41^, residual RRM2 may partially compensate for RRM2B loss. These findings indicate that the extent of mtDNA expansion modulates cGAS–STING activation.

### Eltrombopag inhibits mtDNA release and cGAS-STING dependent inflammation

Having established the RRM2B-cGAS-STING axis, we next sought to pharmacologically block mtDNA release. MtDNA can escape via activation of the mitochondrial permeability transition pore (mPTP), voltage-dependent anion channel 1 (VDAC1), or BAX/BAK megapores^42–44^. We tested eltrombopag (EO), an FDA-approved BAX inhibitor that binds its trigger site and prevents activation^45^.

Immunostaining revealed abundant cytosolic dsDNA in Dox-treated cells, but EO prevented mtDNA release, retaining dsDNA within mitochondria (Fig. 6A-B; Extended Fig. 4A-B). EO also suppressed *CXCL10* but not *IL-8* expression, indicating selective blockade of mtDNA-dependent inflammation (Fig. 6C; Extended Fig. 4C). Comparable effects were obtained with the alternative BAX inhibitor BAI1^46^ (Fig. 6A-C; Extended Fig. 4A-C). Thus, pharmacological inhibition of BAX prevents Dox-induced mtDNA leakage and attenuates RRM2B-driven cGAS–STING signaling in endothelial cells.

**Figure 6.**
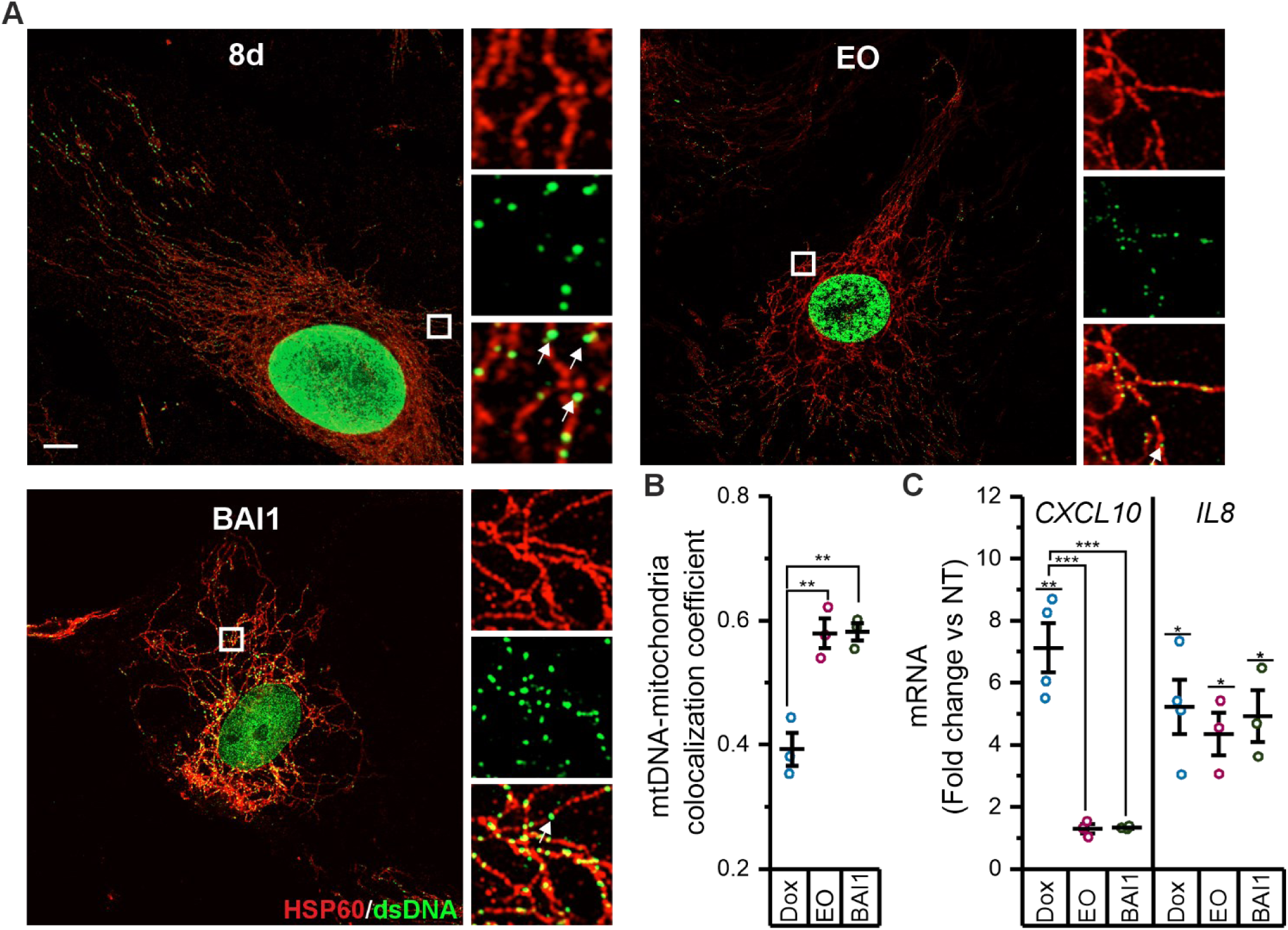
Inhibition of BAX activation prevents mtDNA release and cGAS-STING dependent inflammation. A-B. Immunostaining of HUVECs at 8 days after Dox removal (8d) in the presence or absence of BAX inhibitors, BAX Activation Inhibitor 1 (BAI-1) or eltrombopag (EO), added one day after Dox withdrawal. **A.** Representative super-resolution Airyscan confocal images of dsDNA (green) and HSP60 (red). Scale bar 10 µm. On the right, the magnified images show mtDNA (green) and mitochondria (red), and merge. Arrows indicate cytosolic dsDNA signals. **B.** Quantification of the colocalization coefficient between mtDNA and mitochondria. N=3 independent experiments. ** p<0.01 in a one-way ANOVA test with Tukey’s post hoc analysis for mean comparison. **C.** Expression levels of indicated genes measured by qRT-PCR in HUVECs not treated (NT), and after drug withdrawal (8d) in the presence or absence of BAX inhibitors, BAI-1 or EO. N=3/4 independent experiments. * p<0.05; ** p<0.01; *** p<0.001 in a one-way ANOVA test with Tukey’s post hoc analysis for mean comparison.

### Endothelial RRM2B-STING axis is required for cardiotoxicity *in vivo*

We next examined whether endothelial RRM2B-driven inflammation contributes to DIC *in vivo*. GFP-positive ECs sorted from zebrafish confirmed Dox-induced *rrm2b* upregulation (Extended Fig. 5A), whereas cardiomyocytes showed no increase (Extended Fig. 5B). Endothelial-specific *rrm2b* (rrm2b EC-KD) or *sting* (sting EC-KD) deletion was achieved using *Tg(fli1a:dCas, cryaa:Cerulean)^ia57^* fish^47^ expressing nuclease-deficient Cas9 under the endothelial *fli1a* promoter, and sgRNA microinjection at the one-cell stage (Extended Fig. 6A). qRT-PCR confirmed ∼50% reduction of *rrm2b* or *sting* transcripts in GFP-positive ECs up to 3 dpf, without affecting other tissues (Extended Fig. 6B–E).

In Dox-treated controls, endothelial *rrm2b* induction was accompanied by *ifnphi3*, *irf3*, *il6*, and *il1b* upregulation. In contrast, the increase in *ifnphi3* and *irf3*, but not *il6* or *il1b* was significantly reduced in the *rrm2b* EC-KD fish (Fig. 7A-B), indicating that their induction follows a pathway-specific pattern. Cardiomyocytes from *rrm2b* EC-KD fish also showed diminished *ifnphi3* and *irf3*, despite unaltered *rrm2b* expression (Fig. 7C-D), suggesting paracrine regulation of myocardial inflammation from the endothelium. Similar results were obtained in *sting* EC-KD fish (Extended Fig. 7A-B). Functionally, both *rrm2b* EC-KD and *sting* EC-KD were protected from Dox-induced cardiotoxicity, showing preserved heart rate and reduced cardiac myeloid infiltration (Fig. 7E-G; Extended Fig. 7C-E). These findings establish endothelial *rrm2b* as an upstream regulator of cGAS–STING–mediated inflammation and a key contributor to DIC.

**Figure 7:**
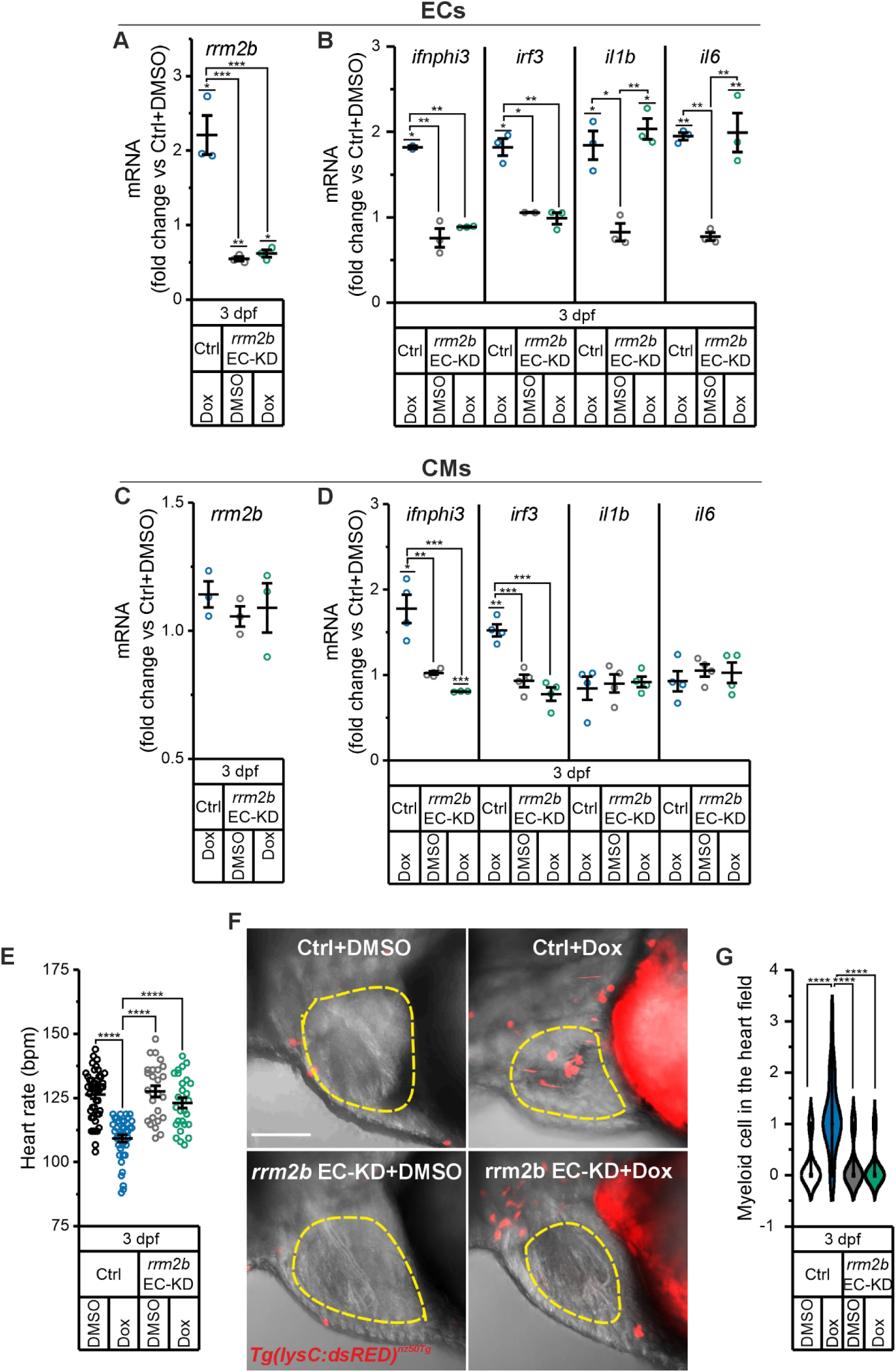
Loss of endothelial *rrm2b* confers protection from doxorubicin cardiotoxicity. **A-B.** Expression levels of **A.** *rrm2b* and **B**. inflammatory genes determined by qRT-PCR in GFP+ ECs sorted from Ctrl and *rrm2b* EC-KD *Tg(kdrl:GFP)^s843Tg^*individuals at 3 dpf with or without Dox treatment. N=3 independent experiments. * p<0.05; ** p<0.01; *** p<0.001 in a one-way ANOVA test with Tukey’s post hoc analysis for mean comparison. **C-D.** Expression levels of **C.** *rrm2b* and **D**. inflammatory genes determined by qRT-PCR in GFP+ CMs sorted from Ctrl and *rrm2b* EC-KD *Tg(tg:EGFP, myl7:EGFP)^ia300Tg^* individuals at 3 dpf with or without Dox treatment. N=3/4 independent experiments. * p<0.05; ** p<0.01; *** p<0.001 in a one-way ANOVA test with Tukey’s post hoc analysis for mean comparison. **F.** Heart rate analysis at 3 dpf in Ctrl and *rrm2b* EC-KD with or without Dox treatment. Bpm, beats per minute. Ctrl+DMSO (n=45), *rrm2b* EC-KD+DMSO (n=38), Ctrl + Dox (n=26), *rrm2b* EC-KD + Dox (n=27). N=3 independent experiments. **** p<0.0001 in a Kruskal-Wallis ANOVA test with Dunn’s post hoc analysis for mean comparison. **G.** Representative confocal image of the heart and myeloid cells in embryos from the transgenic line *Tg(lysC:dsRed2)^nz50Tg^* at 3 dpf. Red dots indicate the dsRED+ myeloid cells, while dotted lines contour the heart. Scale bar 100 µm. **H.** Quantification of myeloid cells in fish heart. Ctrl+DMSO (n=22), *rrm2b* EC-KD+DMSO (n=21), Ctrl + Dox (n=20), *rrm2b* EC-KD + Dox (n=20). N=3 independent experiments. **** p<0.0001 in a Kruskal-Wallis ANOVA test with Dunn’s post hoc analysis for mean comparison.

### Eltrombopag protects against DIC without impairing Dox antitumor efficacy

Since EO blocked RRM2B-driven inflammation by preventing mtDNA release *in vitro*, we tested its cardioprotective efficacy *in vivo*. Embryos were co-treated with Dox and EO and maintained in EO-containing water until 3 dpf, when cardiac dysfunction typically appears. In GFP-sorted ECs, EO selectively prevented induction of *ifnphi3* and *irf3*, without affecting *il1b* or *il6* (Fig. 8A). Functionally, EO preserved heart rate and prevented myeloid accumulation in the heart (Fig. 8B-D), mirroring the protection conferred by H151. EO alone had no cardiac or inflammatory effects. Comparable results were obtained with BAI1 (Extended Fig. 8A-D), supporting mtDNA-release blockade in endothelial cells as a generalizable cardioprotective strategy. To assess whether EO interferes with Dox anticancer efficacy^45^, GFP-labelled human triple-negative breast cancer (MDA-MB-231) cells were xenografted into zebrafish embryos and exposed to Dox for 24 h. EO co-treatment prevented Dox-induced bradycardia and inflammation (Fig. 8E-F) while preserving Dox-mediated tumour reduction (Fig. 8G-H). Similar outcomes were observed with BAI1 (Extended Fig. 8E-H). Altogether, these results demonstrate that pharmacological inhibition of the mtDNA leakage observed in endothelial cells prevents cardiotoxicity without compromising Dox antitumor efficacy, identifying endothelial RRM2B-driven mtDNA release as a tractable therapeutic target in DIC.

**Figure 8.**
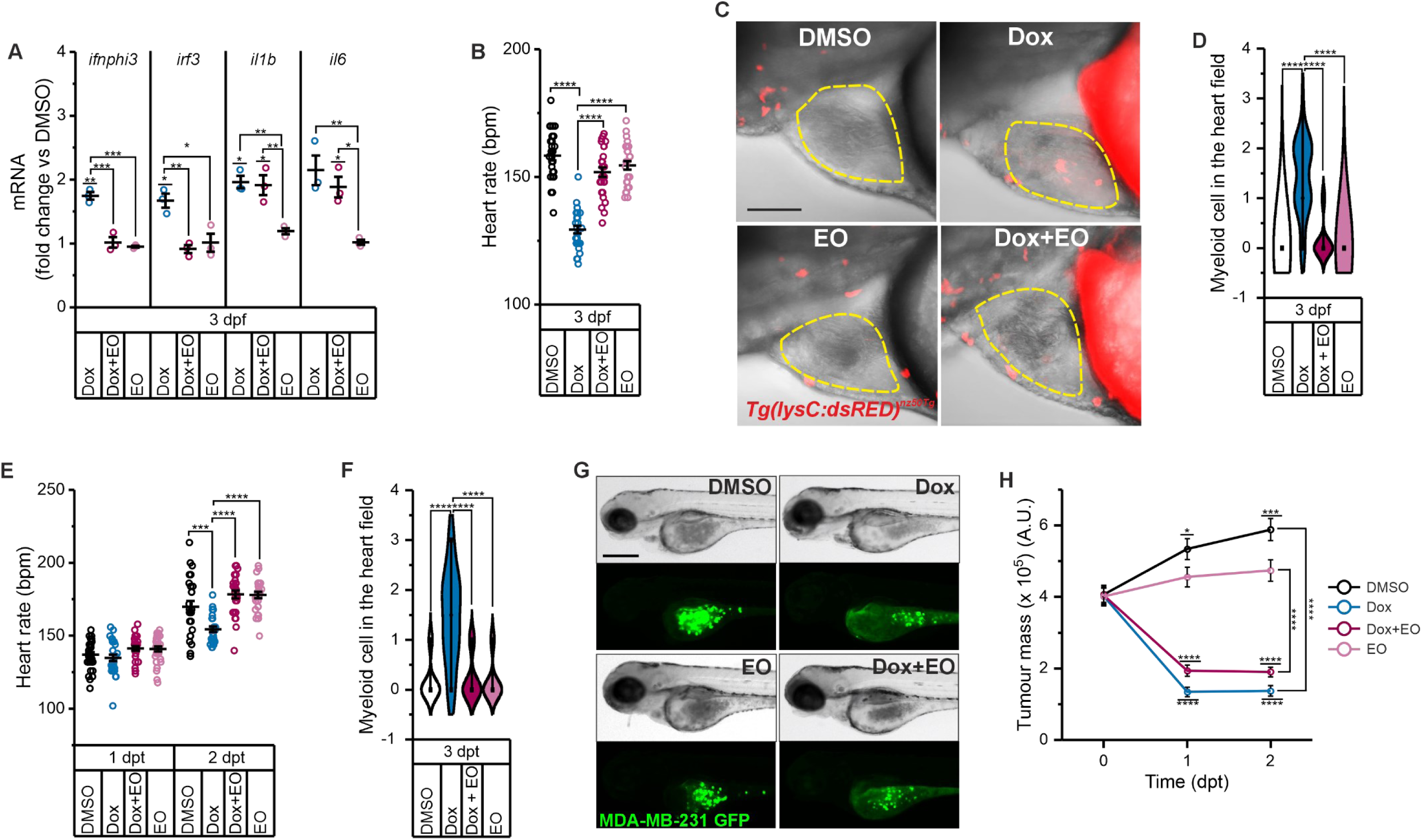
Eltrombopag prevents doxorubicin-induced cardiotoxicity while preserving its antitumor efficacy. **A.** Expression level of the indicated genes in GFP+ sorted ECs from Dox-treated and DMSO control embryos, with or without the BAX inhibitor, EO, of the transgenic line *Tg(kdrl:GFP)^s843Tg^* at 3 dpf. N=3 independent experiments. * p<0.05; ** p<0.01; *** p<0.001 in a one-way ANOVA test with Tukey’s post hoc analysis for mean comparison. **B.** Heart rate analysis in embryos treated as indicated at 3 dpf. Bpm, beats per minute. DMSO (n=27), Dox (n=27), Dox + EO (n=27), EO (n=27). N=3 independent experiments. **** p<0.0001 in a one-way ANOVA test with Tukey’s post hoc analysis for mean comparison. **C-D.** Analysis of myeloid cell recruitment in the heart of embryos from the transgenic line *Tg(lysC:dsRed2)^nz50Tg^*at 3 dpf treated as indicated. **C.** Representative confocal image. Red dots indicate the dsRED+ myeloid cells, while dotted lines contour the heart. Scale bar 100 µm. **D.** Quantification of myeloid cells in fish heart. DMSO (n=25), Dox (n=25), Dox + EO (n=25), EO (n=25). N=3 independent experiments. **** p<0.0001 in a Kruskal-Wallis ANOVA test with Dunn’s post hoc analysis for mean comparison. **E-H.** Human breast cancer (MDA-MB-231) xenograft zebrafish model. Tumour cells were injected at 1 dpf and treated with Dox, or EO or a combination of both drugs (0 dpt, days post treatment). Dox was removed after 24h (1 dpt) whereas EO was maintained in fish water. Analyses were performed at 0, 1 and 2 dpt. **E.** Heart rate analysis. Bpm, beats per minute. DMSO (n=30), Dox (n=30), Dox + EO (n=29), EO (n=30) at 1 dpt; DMSO (n=27), Dox (n=27), Dox + EO (n=27), EO (n=27) at 2dpt. N=3 independent experiments. *** p<0.001; **** p<0.0001 in a Kruskal-Wallis ANOVA test with Dunn’s post hoc analysis for mean comparison. **F.** Quantification of myeloid cells in fish heart of 2 dpt embryos from the transgenic line *Tg(lysC:dsRed2)^nz50Tg^* treated as described. DMSO (n=18), Dox (n=18), Dox + EO (n=18), EO (n=18). N=3 independent experiments. **** p<0.0001 in a Kruskal-Wallis ANOVA test with Dunn’s post hoc analysis for mean comparison. **G.** Representative brightfield and GFP images of the tumour mass in xenografted fish at 2 dpt. **H.** Time curve analysis of the tumour mass in xenografted fish. A.U, arbitrary units. DMSO (n=48), Dox (n=47), Dox + EO (n=42), EO (n=40) at 0 dpt, DMSO (n=42), Dox (n=36), Dox + EO (n=39), EO (n=38) at 1 dpt, DMSO (n=38), Dox (n=32), Dox + EO (n=36), EO (n=36) at 2 dpt. N=3 independent experiments. * p<0.05; *** p<0.001; **** p<0.0001 in a Kruskal-Wallis ANOVA test with Dunn’s post hoc analysis for mean comparison.

## Discussion

Cardiomyocytes and endothelial cells are both a direct target of doxorubicin. Owing to their high mitochondrial content and dependence on oxidative metabolism, cardiomyocytes have long been regarded as the primary site of DIC. In contrast, in ECs mitochondria display a predominant signaling role^48,49^. Whether and how Dox-impaired endothelial mitochondria contribute to DIC, however, has remained unresolved. We identify ECs as active participants in DIC through the p53-inducible ribonucleotide reductase subunit RRM2B, which sustains sterile inflammation and is essential for heart impairment in DIC.

Mitochondrial DNA (mtDNA) is a well-established activator of innate immune signaling^50^, and dysregulated cGAS–STING activation is increasingly recognized as a pathogenic driver in cardiovascular disease^51^. We show that during DIC, endothelial RRM2B induction promotes mtDNA expansion, coupling mitochondrial stress to chronic inflammatory signaling rather than to acute stress responses. Loss-of-function approaches in endothelial cells (mtDNA depletion, genetic inhibition of RRM2B) abolished cGAS–STING activation, identifying mtDNA as the proximal trigger. *In vivo*, the absence of overt cell death, the inflammatory profile observed upon EC specific *sting* or *rrm2b* silencing, and the functional rescue achieved through BAX inhibition collectively implicate mtDNA release as the central event linking Dox exposure to cGAS–STING activation. Notably, RRM2B induction occurred selectively in ECs but not in cardiomyocytes, in line with independent RNA-seq datasets from murine models of DIC^7^, indicating a conserved, endothelial-specific mitochondrial stress program.

In the zebrafish model of DIC established here, vascular alterations preceded overt cardiac dysfunction, mirroring clinical and experimental observations in mammalian systems^52,53^. Given the tight anatomical and metabolic coupling between cardiac ECs and adjacent cardiomyocytes^19^, endothelial inflammatory activation is poised to influence myocardial remodeling. Indeed, endothelial *rrm2b* induction was required for cGAS–STING activation not only within the vasculature but also in neighboring cardiomyocytes, consistent with a paracrine inflammatory circuit. This endothelial response was accompanied by myeloid cell recruitment to the heart, suggesting that both endothelial-derived cytokines and immune-cell infiltration contribute to subsequent myocardial injury^54^. Together, our findings redefine the temporal hierarchy of DIC: endothelial stress arises upstream of cardiomyocyte dysfunction, positioning the vasculature as a trigger rather than a mere bystander of myocardial injury. This concept accords with emerging evidence that ECs orchestrate cardiac remodeling under pathological stress through secreted factors with paracrine and autocrine effects^20^.

Current cardioprotective strategies, including dexrazoxane, provide incomplete protection and may interfere with antitumor efficacy of Dox. Remarkably, pharmacological inhibition of BAX-dependent mtDNA leakage using eltrombopag or BAI1 preserved cardiac function without compromising Dox antineoplastic activity in xenografts *in vivo*. Eltrombopag is FDA approved and its safety in oncology patients with thrombocytopenia is established ^55–57^. Our findings therefore open a potential translational avenue for selective cardioprotection. Targeting mtDNA-driven cGAS–STING activation in endothelial cells may afford mechanistic specificity, mitigating cardiac inflammation without broadly suppressing immune surveillance essential for tumor control. Validation of these mechanisms in mammalian systems, including mouse models of DIC and human iPSC-derived minihearts will further expand the translational safety and scope of eltrombopag across cancer contexts.

In conclusion, we identify endothelial mitochondria-driven sterile inflammation as a determinant of Dox-induced cardiotoxicity. We implicate mtDNA-driven innate immune activation as an initiating event in endothelial–myocardial crosstalk, reframing the pathophysiological sequence of DIC and establishing a foundation for the rational design of next-generation cardioprotective strategies in cancer therapy.

## Methods

### Antibodies and other reagents

The antibodies and reagents were obtained commercially (Supplementary Table 1). All the primer sequences for qRT-PCR, short interference RNAs (siRNAs) used in endothelial cells and sgRNA sequences used in zebrafish are found in Supplementary Table 2.

### Cell lines and treatments

Human endothelial cell lines HUVECs (pooled donors, C2519A, Lonza) and HCMECs (Caucasian male, C-12285 PromoCell) were used between passage 3 and 6. Cells were cultured onto 0.2% −0.5% gelatin-coated plates with Endothelial Cell Growth Medium 2 Kit and Endothelial Cell Growth Medium MV 2 Kit supplemented with 2% and 5% foetal calf serum at 37 °C in a humidified atmosphere and 5% CO_2_ and 5% O_2_.

Mitochondrial DNA-depleted HUVECs (HUVECs ρ⁰) were generated as described^58^. Cells were treated with 50 ng/mL ethidium bromide for 14 days in growth medium supplemented with 5 mM sodium pyruvate and 50 µg/mL uridine. Mitochondrial DNA content/cell (mitochondrial DNA copy number relative to nuclear) was measured by quantitative PCR. Human breast adenocarcinoma cells MDA-MB-231 (HTB26, ATCC) were stably transfected with pEGF-N1. Cells were cultured in DMEM supplemented with 10% FBS and 1% penicillin–streptomycin, and stable expressants were selected by adding 750 µg/ml G-418 (Geneticin) to the medium. All cell cultures were periodically tested for mycoplasma contamination using Venor®GeM Classic mycoplasma detection kit.

HUVECs and HCMECs were exposed to doxorubicin at final concentrations of 250 nM and 200 nM. After 24 hours, the drug-containing medium was removed, cell monolayers were rinsed twice with pre-warmed medium and then maintained in drug-free medium for an additional 8 days, with medium replacement every 2 days. Inhibitors were prepared as stock solutions in DMSO (H151, BAI1, and EO) or nuclease-free water (HU) and added individually on day 2 (one day after Dox removal) at final concentrations of 2 mM (HU), 200 nM (H151), 100 nM (BAI1), and 50 nM (EO).

SiRNA were delivered using Dharmafect-4 achieved following manufacturer instructions. For RRM2B downregulation we used an equimolar combination of Hs_RRM2B siRNA 7 and 8 (30 nM final concentration, Supplementary Table 2), with AllStars Negative Control siRNA (30 nM) as negative control. Three days post-transfection, cells were seeded onto new plates and treated with Dox as described. To sustain RRM2B silencing, cells were transfected again after 5 days.

### Senescent-associated β-Galactosidase Assay

Senescent-associated β-galactosidase (SA-β-gal) was assessed using the Senescent Detection Kit following manufacturer instructions. Cells were fixed in 20% formaldehyde, 2% glutaraldehyde in phosphate-buffered solution for 10 min and incubated overnight at 37°C with X-gal staining solution. SA-β-gal positive cells were visualised under bright-field microscopy using a DMI4000B microscope equipped with Leica HCX PL Fluotar 10x/0.3 M25 objective (Leica). Data are expressed as percentage of SA-β-gal positive cells over imaged cells (>100 cells/sample).

### Immunofluorescence

Cells grown on gelatin-coated plates were fixed in 4% PFA in PBS for 15 min at room temperature, permeabilised with 0.2% (v/v) Triton X-100 in PBS for 10 min at RT, blocked in MAXBlock™ Blocking medium for 1 h at 37°C and rinsed with PBS. Cells were then incubated with antibodies against anti-dsDNA and anti-HSP60 diluted in filtered PBS containing 5% BSA at 4°C overnight, followed by 6 x 5-min washes in 0.05% Tween 20 in PBS (T-PBS).

Secondary antibodies (Alexa Fluor™ 568 donkey anti-rabbit IgG and Alexa Fluor™ 488 donkey anti-mouse) were incubated for 1 h at RT in filtered PBS containing 5% BSA and removed by 4 x 5 min washes T-PBS.

Immunostained cells were then sealed with coverslips using Prolong™ Gold antifade reagent with DAPI. Confocal images were acquired using a LSM 900 microscope in Airyscan mode, equipped with a Plan-Apochromat 63x/1.4 Oil DIC M27 objective (Zeiss). Fluorophores were imaged using sequential line acquisition to prevent spectral bleed-through. Alexa Fluor 488 was excited at 493 nm, with emission at 517 nm, and the detection window was set to 496–565 nm. Alexa Fluor 568 was excited at 577 nm, with emission at 603 nm, and the detection window was set to 577–650 nm. Laser power, detector gain, and pinhole size were kept constant across all conditions. Airyscan images were acquired in super-resolution mode and processed in Zeiss Zen software using standard settings to maximize signal-to-noise and resolution. Image analysis was performed using ImageJ (National Institutes of Health). Mitochondrial length was calculated using the Freehand Line Selection^59^ tool, co-localization between mitochondria and dsDNA using the Squassh plug-in by computing an object-based colocalization coefficient^60^. At least 10 cells per condition were analysed per experiment.

### Mitochondrial DNA copy number quantification

Mitochondrial DNA copy number was determined by real time PCR, as described in.^61^. Genomic DNA was extracted using Puregene Core kit A. Mitochondrial rRNA 12S TaqMan probe and primers rRNA 12S were used to quantify mtDNA. RNase P primers and VIC probe mix were used for nuclear DNA quantification. To quantify mtDNA and nuclear DNA, we used calibration curves generated by serial dilution of a mixture of plasmids carrying the two PCR amplicons. Each DNA sample was analysed in triplicate.

### Detection of mtDNA in the cytosolic fraction

Cytosolic fractions were isolated from control and Dox-treated HUVECs^62^. Cells (1×10^6^) were collected, washed in ice-cold PBS, and resuspended in digitonin lysis buffer (30 µg/ml digitonin in 150 mM NaCl and 50 mM HEPES, pH 7.4) for 10 min at 4°C under gentle end-over-end rotation to permeabilize the plasma membrane selectively. A negative control (NC) was prepared by incubating cells in lysis buffer without digitonin. Samples were centrifuged at 950 × g for 5 min at 4°C to obtain a cytosolic supernatant fraction and a pellet of permeabilized cells. The supernatant was transferred into new tubes and clarified by centrifugation at 17,000 × g for 5 min at 4°C. The pellet was discarded, and the supernatants were collected as the cytosolic fractions. The pellet was washed three times with ice-cold PBS to remove cytosolic contamination and lysed in SDS lysis buffer (1% SDS, 20 mM Tris-HCl, pH 8) at 95°C for 15 min. After clarification at 17,000 × g for 5 min at 4°C, the supernatant was collected as mitochondria-nuclear fraction. One-fifth of the cytosolic and mitochondria-nuclear fractions were used for protein analysis Samples were boiled at 95°C for 5 min, vortexed, and clarified at 17,000 × g for 5 min. Equal amounts were resolved by SDS–PAGE and probed with antibodies against each cellular compartment. Vinculin, TOM20 and HIS H3 were used as a cytosolic, mitochondrial and nuclear marker. DNA was isolated from cytosolic fractions by Phenol Chloroform-based method, after incubation with 50 mg/ml of RNase A for 1 h at 37°C, followed by 1 h at 55 °C with 2 mg/ml Proteinase K Solution. qPCR was performed on using primers for mtDNA (12S rRNA) and nDNA (RNase P), as previously described. The cytosolic mtDNA copy number per 10^6^ cells was calculated by normalising the cytosolic mtDNA copies of each sample (S) to that of the corresponding negative control (NC), according to the following formula:

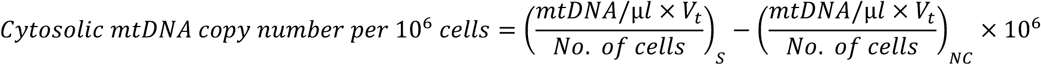

where *mtDNA copies/µl* represents the cytosolic mtDNA copy number determined by qPCR in 1 µl of the fraction, *V_t_* is the total DNA volume of the *cytosolic fraction*, and *No. of cells* corresponds to the number of cells lysed for the experiment.

### BrdU pulse and chase assay

mtDNA synthesis and degradation were assessed by pulse and chase experiments with 5-bromo-2′-deoxyuridine (BrdU) followed by immunofluorescence. Dox-treated HUVECs at day 8 were incubated for 4 and 8 hours with 10 μM BrdU. In chase experiments, we removed the BrdU-medium after 8 hours, rinsed the monolayer twice with prewarm medium and incubated the cells for additional 16 hours in BrdU-free medium. After the incubation, cells were fixed with 50 mM glycine pH 2.0 in 70% ethanol for 35 min at −20°C and BrdU incorporated into the newly synthesized mtDNA was detected using the BrdU Labeling and Detection kit I according to the manufacturer’s instructions. Nuclei were counterstained using Prolong™ Gold Antifade Reagent with DAPI. Images were acquired with Zeiss LSM 900 microscope (Zeiss), a Plan-Apochromat 63x/1.4 Oil DIC M27 objective. Fluorophores were imaged using sequential line acquisition to prevent spectral bleed-through. DAPI was excited with the 405 nm diode laser and emission was collected at 420–480 nm. Alexa Fluor 488 was excited at 493 nm, with emission at 517 nm, and the detection window was set to 496–565 nm. Laser power, detector gain, and pinhole size were kept constant across all conditions. Airyscan images were acquired in super-resolution mode and processed in Zeiss Zen software using standard settings to maximize signal-to-noise and resolution. Image analysis was performed using Analyze particle tool in ImageJ (National Institutes of Health). Data were expressed as the number of BrdU puncta per cell, with a total of 40 cells analysed per time point.

### Total RNA extraction and qRT–PCR

To quantify messenger RNA transcript abundance, total cellular RNA was extracted using TRIzol Reagent according to the manufacturer’s protocol. RNA concentration and purity were determined using NanoDrop Lite Spectrophotometer (Thermo Fisher Scientific) at A260 nm and A280/260, respectively. cDNA was synthesised from 1 µg of total RNA using the High-Capacity cDNA Reverse Transcription Kit according to the manufacturer’s instructions. Equal amount of cDNA was used for qRT-PCR using HOT FirePol EvaGreen qPCR Supermix 5× and the appropriate primers (Supplementary table 2) on QIAquant 384 5plex (Qiagen). For each biological samples, three technical replicates were performed and normalized against the housekeeping genes human *alpha Tubulin* (*TUBA)* or zebrafish *elongation factor 1-alpha 1* (*eef1a1*). Relative expression was analysed using the 2^−ΔΔCt^ method, and the relative fold change was plotted with the control samples given a value of 1. Results are presented as mean ± SEM.

### SDS-PAGE and immunoblot analysis

Cells were collected, washed with cold PBS and lysed on ice for 30 minutes in RIPA buffer supplemented with 1x phosphatase inhibitor and 1x protease inhibitor cocktail. The soluble lysate fractions were clarified by centrifugation at 19,000 × g for 15 min, transferred in new tubes and stored at −80°C. Protein concentrations were determined by Bradford assay. Equal amounts of proteins (15–20 μg) were resolved using a 4-12% SDS-PAGE Gel, followed by transfer to a nitrocellulose membrane. The membrane was blocked with 4% milk in Tris-buffered saline with 0.1% Tween 20 (TBS-T) for 1 h at RT, followed by immunoblotting with antibodies listed in Supplementary Table 1. The blot was developed using Immobilon^®^ Forte or Classico Western HRP Substrates and imaged with the chemiluminescence imaging analyzer iBright 1500 Imaging System (Thermo Fisher Scientific). Band intensities were quantified by ImageJ software (National Institutes of Health). Each sample was normalized on the housekeeping protein and reported as fold change relative to control sample given as value 1.

### Maintenance and handling of zebrafish transgenic line

All *Danio rerio* (zebrafish) experiments were performed in accordance with the Italian and European Legislations (Directive 2010/63/EU) and with permission for animal experimentation from the Ethics Committee of the University of Padua and the Italian Ministry of Health (Authorization number 728/2025-PR).

The experiments were performed on embryos and larvae up to 5 days post-fertilisation (dpf), before the onset of independent feeding. Zebrafish were maintained in a temperature-controlled environment at 28.5°C, with a 12:12 light/dark cycle, and fed as described by Kimmel *et al*.^63^. For the anaesthesia of zebrafish embryos and larvae, buffered Tricaine was added to their water at 0.16 mg/ml. Wild-type lines used in this work included the Tuebingen, Giotto, and Umbria strains. For *in vivo* studies, the following transgenic lines were used: *Tg(kdrl:GFP)^s843Tg^*, *Tg(lysC:dsRed2)^nz50Tg^*, *Tg(tg:EGFP, myl7:EGFP)^ia300Tg^*, *Tg(fli1a:dCas9, cryaa:Cerulean)^ia57^*.

### Chemical treatments *in vivo*

Zebrafish embryos were manually dechorionated at 1 dpf and exposed to Dox (10-30 μM) or 0.1% DMSO in E3 medium supplemented with 0.003% PTU to inhibit pigmentation. After 24 hours, Dox was washed out, and larvae were transferred to fresh PTU-containing E3 medium and kept until 5 dpf. Inhibitors H151, BAI1 and EO were co-administered with Dox from 1 dpf and maintained in the medium throughout the experimental end point at the final concentration of 2 µM (H151), 5 µM (BAI1) and 1 µM (EO).

### Survival analysis

Following Dox treatment embryo survival was monitored up to 5 dpf. Embryos were scored as *dead* based on the absence of heartbeat and spontaneous movement. Survival data were analysed using the Kaplan-Meier method, and the survival curves were compared using the log-rank test. Statistical analyses were performed using OriginPro (Version 2025, OriginLab Corporation), with *p* < 0.05 considered statistically significant.

### Evaluation of zebrafish cardiac function

Upon Dox cardiac function was assessed at 2, 3, and 5 dpf by measuring heart rate, diastolic atrial area, fractional shortening (FS), and pericardial oedema^64,65^. Heart rate was determined manually by counting cardiac contractions over a 15-second interval in anaesthetised larvae using a stopwatch. Counts were performed in triplicate for each individual, and the average was multiplied by four to obtain beats per minute (bpm).

Diastolic atrial area and fraction shortening were assessed using the *Tg(tg:EGFP, myl7:EGFP)^ia300Tg^* transgenic zebrafish line. Anesthetised larvae were embedded in 2% methylcellulose, and images and videos of contracting hearts were acquired using a Leica M165FC stereomicroscope equipped with a DFC7000T digital camera (Leica). Video analysis was performed using ImageJ software (National Institutes of Health). Diastolic atrial area was measured at the end of atrial diastole using the Freehand selection tool, averaged across three cardiac cycles per larva, and normalised to body length. FS was calculated by measuring the ventricular end-diastolic diameter (EDD) and end-systolic diameter (ESD) using Straight Line tool, applying the formula^66^:

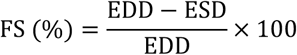

Pericardial oedema was assessed by visual inspection under a stereomicroscope. Larvae were scored for the presence or absence of visibly enlarged pericardial chambers, and the percentage of affected larvae was calculated by dividing the number of larvae showing oedema by the total number of individuals.

### Image acquisition and analysis of zebrafish larvae

*Vascular morphology* was evaluated by measuring the diameter of the dorsal aorta at the level of 8^th^ somite in *Tg(kdrl:GFP)^s843T^* transgenic line zebrafish embryos and larvae. Anaesthetised larvae were embedded in 2% low-melting point agarose. Confocal images were acquired using a Nikon C2 confocal microscope equipped with Plan Fluor 20X/0.50 DIC M/N2 ∞/0.17 WD 2.1 objective (Nikon). Image analysis was performed using ImageJ software (National Institutes of Health) and the dorsal aorta diameter was measured using the Straight-Line Selection tool.

Myeloid cells in the heart were visualized using the transgenic zebrafish line *Tg(lysC:dsRed2)^nz50Tg^*. Larvae were anaesthetised and mounted in 2% low-melting point agarose. Z-stack images of the heart were acquired using a Nikon C2 confocal microscope equipped with Plan Fluor 20X/0.50 DIC M/N2 ∞/0.17 WD 2.1 objective (Nikon). The number of dsRED-positive myeloid cells in the heart was manually quantified across the z-stack.

### Zebrafish embryos dissociation for fluorescent-activated cell sorting

GFP-positive ECs and CMs were isolated from larvae of the transgenic lines *Tg(kdrl:GFP)^s843Tg^* and *Tg(tg:EGFP, myl7:EGFP)^ia300Tg^* by fluorescent-activated cell sorting (FACS) as reported in Manoli M *et al*. ^67^. Briefly, pools of 50-100 embryos were dissociated at different dpf in PBS containing 0.25% phenol red–free trypsin, 1 mM EDTA (pH 8.0), and 2.2 mg/mL collagenase type II. Digestion was quenched with 1 mM CaCl₂ and 10% FBS. The resulting cell suspension was washed in PBS, resuspended in Opti-MEM supplemented with 10% FBS and 1× penicillin–streptomycin, and passed through a 40-μm nylon mesh. Cell sorting was performed using a FACSAria III cell sorter (BD Biosciences) equipped with blue (488 nm), red, violet, and yellow-green lasers. EGFP-positive cells were selected based on forward scatter (FSC) versus side scatter (SSC) profiles and sorted using a 20 mW 488 nm laser with 502 LP and 530/30 BP filters. Data acquisition and analysis were performed using FACSDiva software (BD Biosciences). Sorted populations consistently achieved ≥95% purity. Cells were collected in resuspension medium and processed for RNA isolation using the NucleoSpin RNA XS kit, following the manufacturer’s protocol.

### Annexin V staining

Cell death was assessed by annexin V binding assay in GFP-positive ECs and CMs isolated from 2 dpf zebrafish embryos of the transgenic lines *Tg(kdrl:GFP)^s843Tg^* and *Tg(tg:EGFP, myl7:EGFP)^ia300Tg^*, treated with 10 or 30 μM Dox for 24 h. Although the two concentrations are displayed separately in the figures for clarity, experiments with 10 and 30 μM Dox were performed simultaneously under identical experimental conditions. Pools of 50-100 embryos were dissociated in 1× PBS containing 0.25% trypsin phenol red-free, 1 mM EDTA (pH 8.0) and 2.2 mg/ml collagenase type II. Digestion was quenched with 1 mM CaCl₂ and 10% FBS. The resulting cell suspension was washed in PBS, resuspended in Opti-MEM supplemented with 10% FBS and 1× penicillin–streptomycin, and passed through a 40-μm nylon mesh. Annexin V staining was performed using the Annexin V Apoptosis Detection Kit with Annexin V PE-Cy7, following the manufacturer’s instructions. The percentage of Annexin V-positive cells within the GFP-positive populations was quantified by flow cytometry (FACSCanto II, BD Biosciences), and data were analysed using BD FACSDiva™ Software (BD Biosciences).

### CRISPRi-mediated endothelial-specific gene downregulation in zebrafish

Targeted downregulation of *rrm2b* and *sting* in endothelial cells was achieved using CRISPR interference (CRISPRi) technology. sgRNAs were designed with the Breaking-Cas tool^68^ to target exons 3 and 4 of either *rrm2b* or *sting*. Chemically modified sgRNAs were synthesised by Synthego (Synthego Corporation) and resuspended in nuclease-free water. A mix of two sgRNAs was microinjected into one-cell stage embryos of the transgenic zebrafish line *Tg(fli1a:dCas9, cryaa:Cerulean)^ia57^* ^47^ using a WPI PicoPump microinjector (Sarasota). The sgRNA sequences targeting *rrm2b* and *sting* were reported in Supplementary Table 2. Downregulation of *rrm2b* and *sting* was confirmed by qRT-PCR in FACS-sorted endothelial cells. Embryos were then treated with Dox as previously described, and both cardiac function and inflammatory responses were assessed at 3 dpf.

### Cancer cell xenotransplant

For cancer cell xenotransplantation, 1 dpf embryos were manually dechorionated and anaesthetised. 1 x 10^5^ cells/µl MDA-MB-231-GFP cells were injected into the yolk as a single droplet (around 100 cells per embryo) using a WPI PicoPump microinjector (Sarasota). After 5 hours, the embryos were fluorescently assessed for successful cell implantation and subjected to drug treatment with 10 µM Dox, 1 µM EO, 5 µM BAI1, and their combinations. Although treatment groups are displayed separately in the figures for clarity, all drug conditions were initiated and conducted simultaneously under identical experimental settings. Xenotransplanted embryos were maintained at 33°C, and viability was monitored daily. At 1 and 2 days post treatment (dpt) larvae were anaesthetised, and images of the tumours were acquired using AXIO Zoom V16 stereomicroscope equipped with an Axiocam 305 Mono camera with 30.5x magnification (Zeiss). Tumour mass was quantified using ImageJ (NIH, Bethesda, MD, USA) by calculating the integrated density of GFP-expressing MDA-MB-231 cells within a predefined region of interest (ROI), following subtraction of background signal.

### Statistical analysis

Statistical analyses were performed in OriginPro (Version 2025, OriginLab Corporation). All data were based on independent biological replicates. Independent cell culture experiments were performed by thawing fresh aliquots from the original cell stock.

Normality of the data was confirmed using the Shapiro-Wilk test. If the data passed the normality test (a = 0.05), then a parametric test, such as unpaired t-test or ordinary one-way ANOVA was used. If the data did not pass the normality test, the nonparametric tests (Mann–Whitney U-test or Kruskal–Wallis test for multiple comparisons) was used. Multiple comparisons were corrected using Tukey’s or Dunn’s test, as appropriate.

Data are presented as dot plot showing individual data points, mean, and SEM, from at least 3 independent biological replicates, including the exact *P* values, with *P* < 0.05 considered statistically significant. The individual statistical tests used, as well as the exact value of sample size (n, indicates the number of animals/samples) for *in vivo/in vitro* experiments, are mentioned in the corresponding figure legends.

## Acknowledgement

The authors would like to thank Ms. S. Iljazi and the staff of the Zebrafish Facility at DeBio, University of Padova, for their valuable technical assistance, Dr. A. Cabrelle of the Veneto Institute of Molecular Medicine for technical support with FACS sorting and cytofluorimetric analyses. We are grateful to Prof. Robert N. Wilkinson for kindly providing the plasmid used to generate the transgenic line *Tg(fli1a:dCas;cryaa:cerulean)^ia57^* employed in this study. This study was supported by European Research Council ERC GA282280; Ministero dell’Università e della Ricerca PRIN2020PKLEPN_002 (L.S.); European Union-Next Generation EU, Mission 4, Component 2, CUP B93D21010860004 (L.S.); Ministero dell’Università e della Ricerca PRIN2022PNRR P2022JZ9RE (L.S.); S.S. is the recipient of the SOE_0000181, MUR Concession Decree No. n. 564 of 13/12/2022, CUP C93C22007650006, funded under the National Recovery and Resilience Plan (NRRP), Mission 4, Component 2, Investment 1.2, MUR Call for tender n. 367 of 7/10/2022 funded by the European Union – NextGenerationEU. M. Z. was the recipient of an AIRC postdoctoral fellowship. R.B.C. was a Post-Doc fellow supported by DeBio, UniPD (grant 2024DIBIO1SIDASSEGNI-00095). G.R. was a Post-Doc fellow supported by the Italian Ministry of University and Research, MUR (grant PRIN 2022WZCXRZ). N.T. was supported by the Italian Telethon Foundation (grant GGP19287) and MUR (grant PNRR M4C2 CN00000041).

## Author contributions

G.P. conceived and supervised the project. S.G. and G.P. designed the experiments, discussed the data, and wrote the manuscript. S.G. and A.B. performed most of the experiments, analysed and interpreted data, and prepared the figures. N.F. generated the transgenic line *Tg(fli1a: dCas9, cryaa:Cerulean)^ia57^*, contributed to the design of zebrafish experiments, and participated in data discussion. R.B.C. and G.R. contributed to zebrafish experiments and data analysis. C.R. performed the mtDNA content analyses. M.Z. and S.S. conducted the immunofluorescence analyses. N.T. performed the xenograft tumour injections and, together with F.A., provided materials. L.S. provided materials and the graphic design tool, participated in data discussion, and offered critical comments on the manuscript. All authors revised the manuscript.

## EXTENDED FIGURES

**Extended Figure 1.**
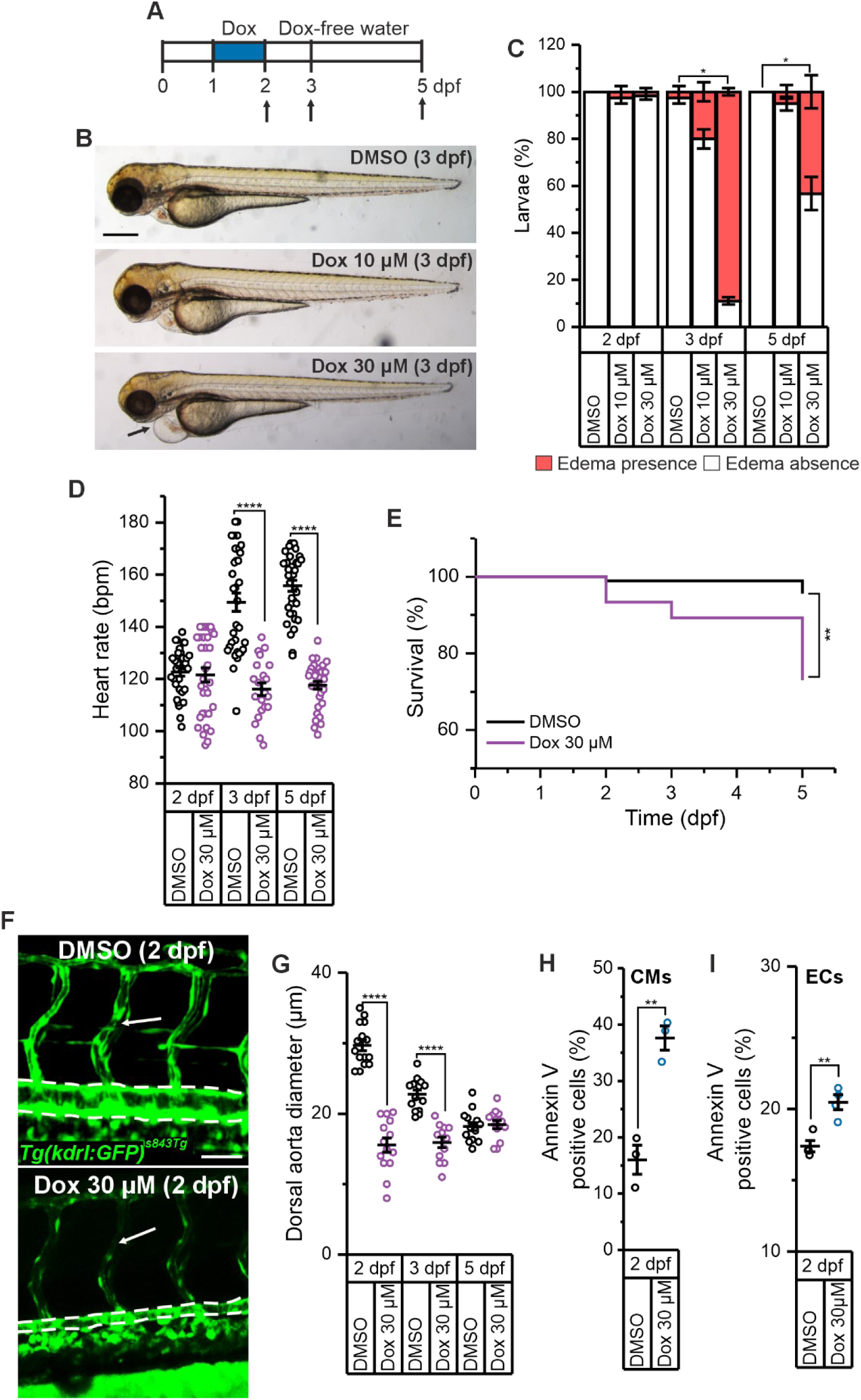
Zebrafish model faithfully recapitulates doxorubicin-induced cardiotoxicity. **A.** Experimental model of Dox-induced cardiotoxicity with timeline of the analysis. Briefly, zebrafish embryos were treated with 10-30 µM Doxorubicin at 1 dpf. After Dox withdrawal at 2 dpf, embryos and larvae were maintained in drug-free water up to 5 dpf. Dox, Doxorubicin-treated fish. DMSO, dimethyl sulfoxide, control fish. Arrows indicate the time of analysis. **B.** Representative images of pericardial oedema analysis in Dox-treated embryos and DMSO controls at 3 dpf, with arrow indicating the pericardial oedema. Scale bar 500 µm. **C.** Pericardial oedema analysis in Dox-treated embryos and DMSO controls. Single individuals were scored for the presence or absence of pericardial oedema. DMSO (n=30), 10 μM Dox (n=30), 30 μM Dox (n=30) at 2 dpf, DMSO (n=29), 10 μM Dox (n=29), 30 μM Dox (n=27) at 3 dpf; DMSO (n=29), 10 μM Dox (n=29), 30 μM Dox (n=25) at 5 dpf. N=3 independent experiments. * p<0.05 in a Kruskal-Wallis ANOVA test with Dunn’s post hoc analysis for mean comparison. **D.** Heart rate analysis at different dpf in Dox-treated embryos and DMSO controls. Bpm, beats per minute. DMSO (n=34), 30 μM Dox (n=35) at 2 dpf, DMSO (n=33), 30 μM Dox (n=21) at 3 dpf, DMSO (n=35), 30 μM Dox (n=34) at 5 dpf. N=3 independent experiments. **** p<0.0001 in a Mann-Whitney test. **E.** Kaplan-Meier Survival analysis curves of Dox-treated embryos and DMSO controls. DMSO (n=29), 30 μM Dox (n=30) at 2 dpf, DMSO (n=30), 30 μM Dox (n=27) at 3 dpf, DMSO (n=29), 30 μM Dox (n=24) at 5 dpf. N=3 independent experiments. ** p<0.01 in a log-rank test. **F.** Representative confocal images of the trunk vasculature in Dox-treated and control 2 dpf *Tg(kdrl:GFP)^s843Tg^* embryos. Arrows indicate the intersegmental vessels (ISVs), dotted lines contour the dorsal aorta (DA). Scale bar 50 µm. **G.** Quantification of DA diameter in experiments as in F. DMSO (n=15), 30 μM Dox (n=14) at 2 dpf, DMSO (n=14), 30 μM Dox (n=12) at 3 dpf, DMSO (n=14), 30 μM Dox (n=13) at 5 dpf. N=3 independent experiments. **** p<0.0001 in a two-sample t-test with Welch’s post hoc analysis for mean comparison. **H-I.** Average ± SEM of Annexin-V+ of GFP+ cardiomyocytes, CMs (H), and endothelial cells, ECs (I) isolated from 2 dpf *Tg(tg:EGFP, myl7:EGFP)^ia300Tg^* and *Tg(kdrl:GFP)^s843Tg^* embryos treated with DMSO or 30 μM Dox. N=3-4 independent experiments. ** p<0.01 in a two-sample t-test with Welch’s post hoc analysis for mean comparison.

**Extended Figure 2.**
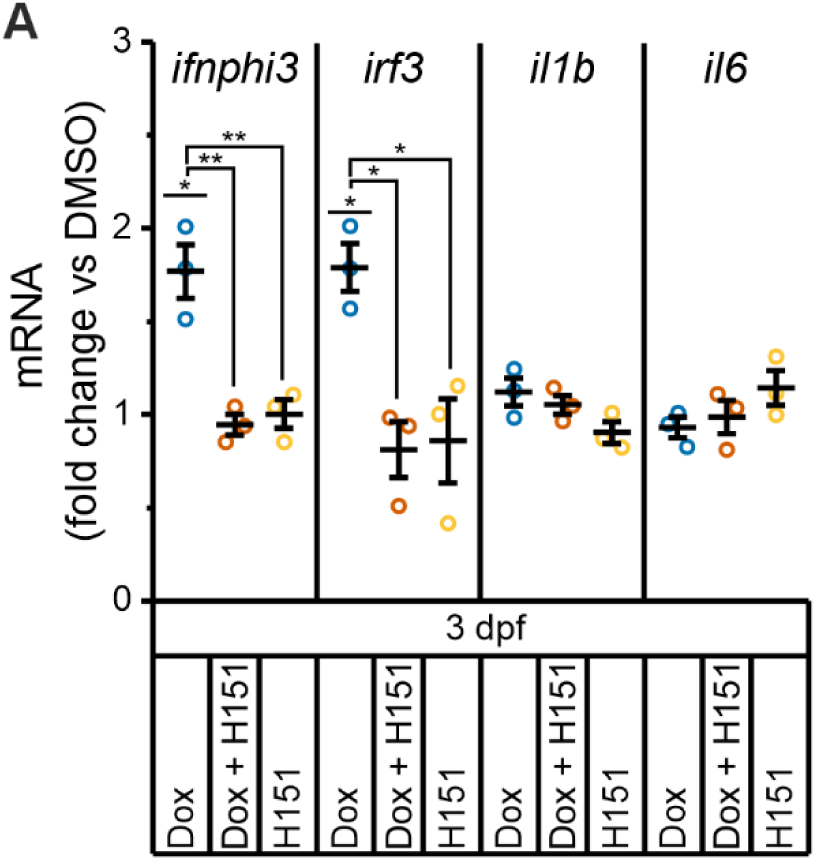
Doxorubicin-induced inflammatory pattern in cardiomyocytes. **A.** Expression level of the indicated inflammatory genes in GFP+ sorted CMs of Dox-treated and DMSO control embryos in the presence or absence of H151 at 3 dpf from the transgenic line *Tg(tg:EGFP, myl7:EGFP)^ia300Tg^*. N=3 independent experiments. * p<0.05; ** p<0.01 in a one-way ANOVA test with Tukey’s post hoc analysis for mean comparison.

**Extended Figure 3.**
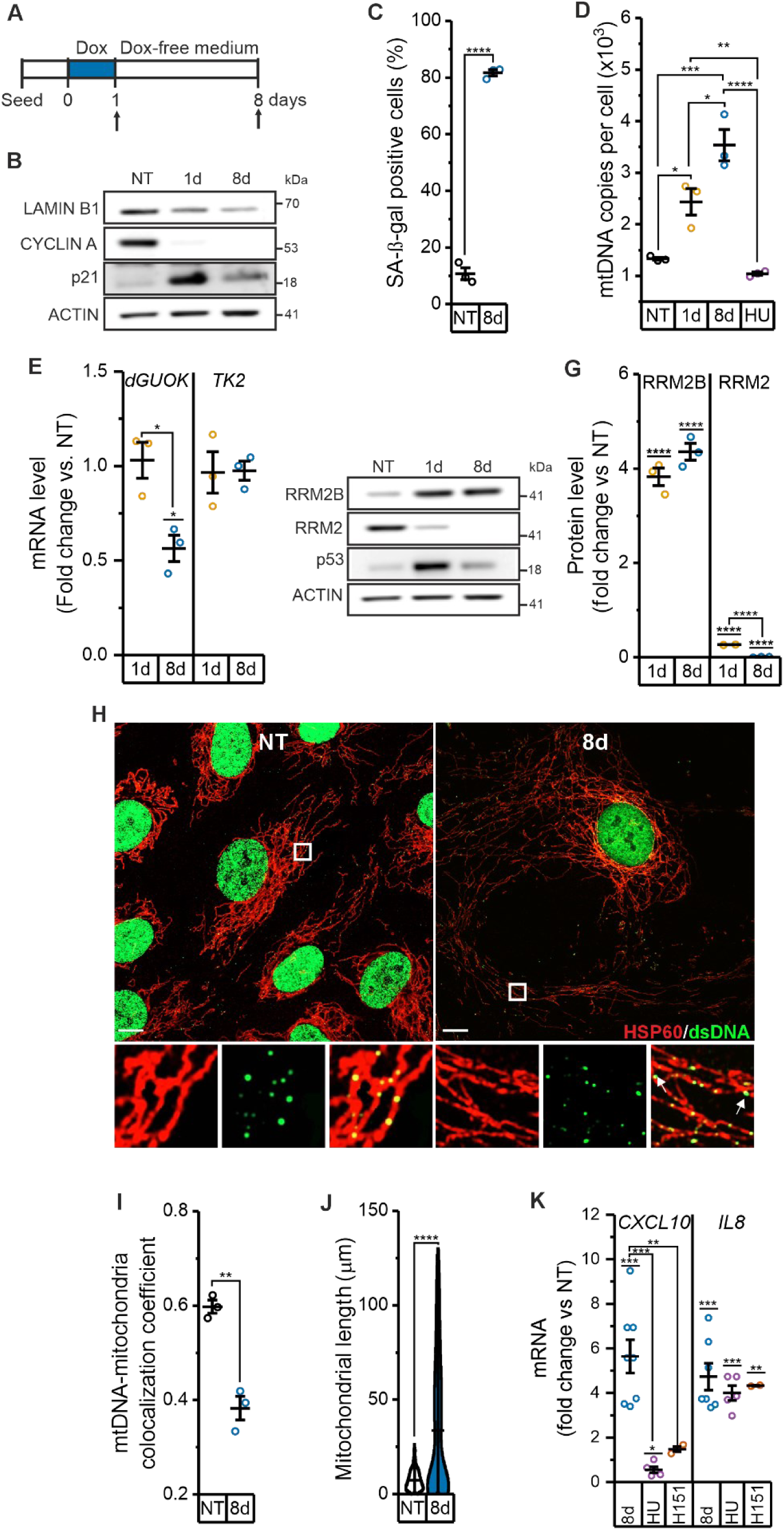
Consequences of Doxorubicin exposure in cardiac endothelial cells *in vitro*: senescence, mtDNA instability and inflammation. **A.** Experimental model of endothelial cells transiently exposed to Doxorubicin^28^. **B.** Representative immunoblot image in HCMECs, not treated (NT), during Dox exposure (1d) and after drug withdrawal (8d). **C.** SA-β-galactosidase assay of HCMECs, not treated (NT), during Dox exposure (1d) and after drug withdrawal (8d). **** p<0.0001 in a two-sample t-test with Welch’s post hoc analysis for mean comparison. **D.** Quantification of mtDNA copy per cell determined by qPCR in HCMECs, not treated (NT), during Dox exposure (1d) and after drug withdrawal (8d). Hydroxyurea (HU) was added one day after Dox withdrawal. N=3 independent experiments. * p<0.05; ** p<0.01; *** p<0.001; **** p<0.0001 in a one-way ANOVA test with Tukey’s post hoc analysis for mean comparison. **E.** Expression level of dGUOK and TK2 by qRT-PCR in HCMECs, during Dox exposure (1d) and after drug withdrawal (8d). TK2, thymidine kinase 2. dGUOK, deoxyguanosine kinase. N=3 independent experiments. * p<0.05 in a one-way ANOVA test with Tukey’s post hoc analysis for mean comparison. **F.** Representative immunoblot image and **G.** quantification of the canonical RRM2 and alternative p53 inducible RRM2B small RNR subunit in HCMECs, not treated (NT), during Dox exposure (1d) and after drug withdrawal (8d). N=3 independent experiments. **** p<0.0001 in in a one-way ANOVA test with Tukey’s post hoc analysis for mean comparison. H-J. Immunostaining of HCMECs not treated (NT) and after Dox withdrawal (8d). **H.** Representative super-resolution Airyscan confocal images of dsDNA (green) and HSP60 (red) Scale bar 10 µm. At bottom, the magnified images show mtDNA (green) and mitochondria (red), showing that most DNA foci are located within mitochondria with some foci in the cytoplasm of Dox treated HCMECs. Arrows indicate cytosolic dsDNA signals. **I.** Quantification of the colocalization coefficient between mtDNA and mitochondria. N=3 independent experiments. ** p<0.01 in a two-sample t-test with Welch’s post hoc analysis for mean comparison. **J.** Quantification of mitochondrial length. NT (n=317), 8d (n=379). N=3 independent experiments. **** p<0.0001 in a Mann-Whitney test. **K.** Expression levels of indicated genes measured by qRT-PCR in HCMECs, not treated (NT) and after drug withdrawal (8d). Hydroxyurea (HU) or Sting inhibitor H151 (H151) was added one day after Dox withdrawal. N=3 independent experiments. * p<0.05; ** p<0.01; *** p<0.001 in a one-way ANOVA test with Tukey’s post hoc analysis for mean comparison.

**Extended Figure 4.**
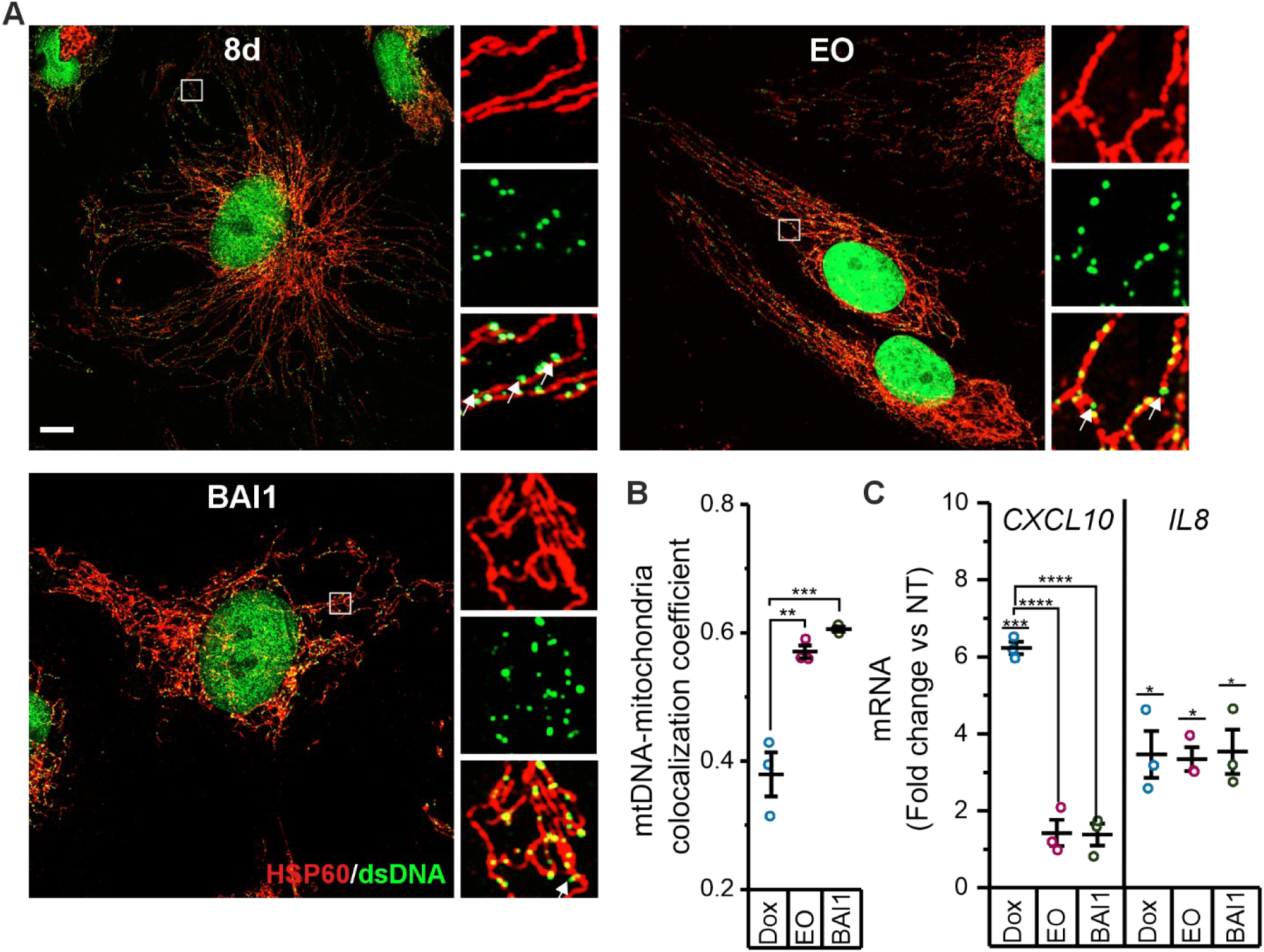
Inhibition of BAX activation prevents mtDNA release and cGAS-STING dependent inflammation. **A-B.** Immunostaining of HCMECs 8 days after Dox removal (8d) in the presence or absence of BAX inhibitors, BAX Activation Inhibitor 1 (BAI1) or eltrombopag (EO), added one day after Dox withdrawal. **A.** Representative super-resolution Airyscan confocal images of dsDNA (green) and HSP60 (red). Scale bar 10 µm. On the right, the magnified images show mtDNA (green) and mitochondria (red), and merge. Arrows indicate cytosolic dsDNA signals. **B.** Quantification of the colocalization coefficient between mtDNA and mitochondria. N=3 independent experiments. ** p<0.01; *** p<0.001 in a one-way ANOVA test with Tukey’s post hoc analysis for mean comparison. **C.** Expression levels of indicated genes measured by qRT-PCR in HCMECs not treated (NT), and after drug withdrawal (8d) in the presence or absence of BAX inhibitors, BAI1 or EO. N=3 independent experiments. * p<0.05; *** p<0.001; **** p<0.0001 in a one-way ANOVA test with Tukey’s post hoc analysis for mean comparison.

**Extended Figure 5.**
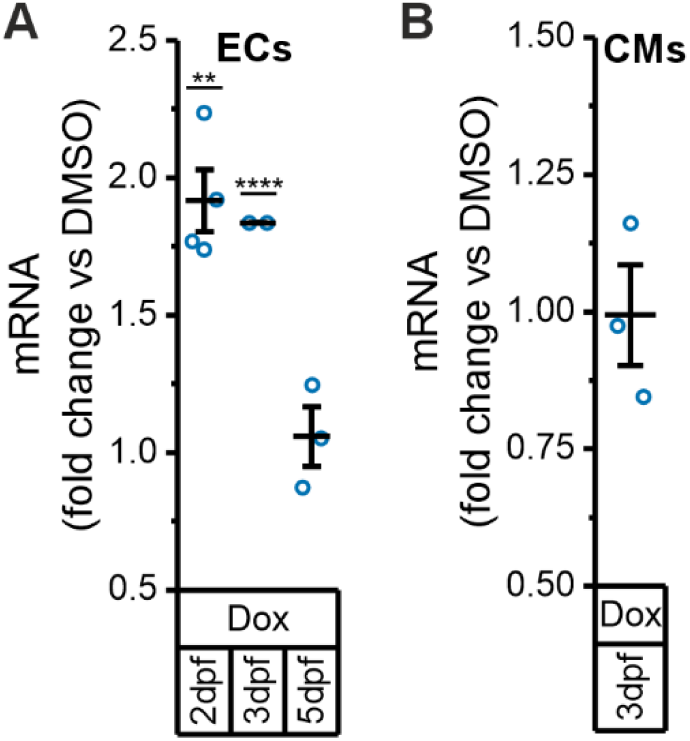
Tissue-specific rrm2b expression upon Doxorubicin exposure. **A.** Expression level of *rrm2b* at the indicted dpf determined by qRT-PCR in GFP+ ECs sorted from *Tg(kdrl:GFP)^s84Tg3^* embryos treated as indicated. N=3 independent experiments. ** p<0.01; **** p<0.0001 in a two-sample t-test with Welch’s post hoc analysis for mean comparison. **B.** Expression level of *rrm2b* determined by qRT-PCR in GFP+ CMs sorted from *Tg(tg:EGFP, myl7:EGFP)^ia300Tg^* embryos at 3 dpf treated as indicated. N=3 independent experiments. p≥ 0.05 in a two-sample t-test with Welch’s post hoc analysis for mean comparison.

**Extended Figure 6.**
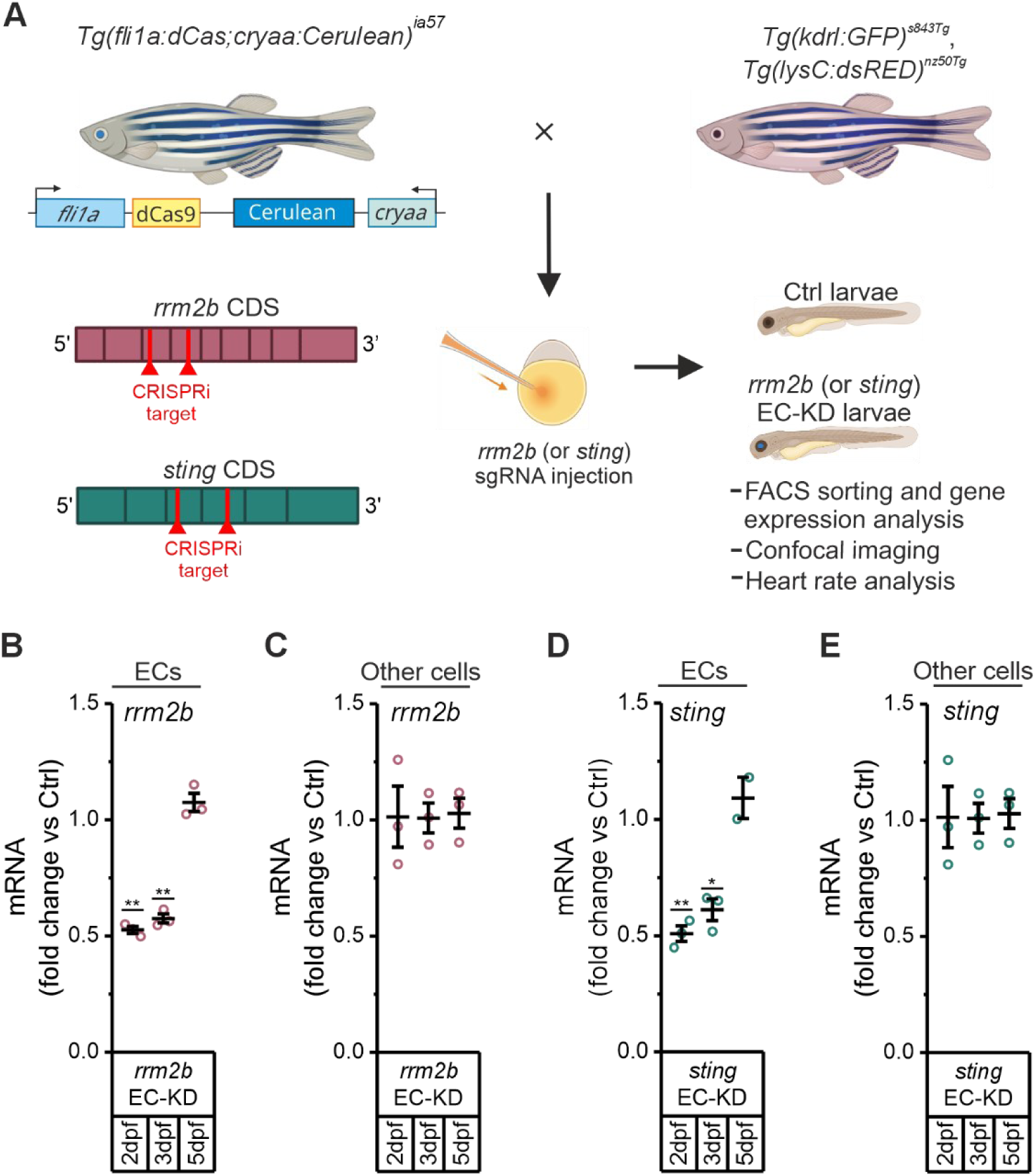
Zebrafish CRISPRi model for endothelial-specific knockdown of *rrm2b* or *sting*. **A.** Experimental model used for endothelial-specific downregulation of *rrm2b* or *sting* in zebrafish. *Tg(fli1a:dCas9, cryaa:Cerulean)^ia59^* zebrafish, expressing a deadCas9 (dCas9) in the endothelium and a Cerulean eye marker for transgenesis tracking, were crossed with the indicated transgenic lines. One-cell stage embryos were injected with sgRNAs targeting exons 3-4 of rrm2b or sting. Cerulean-positive larvae were selected as endothelial-specific knockdown (*rrm2b EC-KD or sting EC-KD*), while Cerulean-negative siblings served as controls (Ctrl). Illustration created with BioRender. **B-C.** Expression level of *rrm2b* in **B.** GFP+ ECs and **C.** GFP-cells (Other cells) sorted from Ctrl and EC-KD *Tg(kdrl:GFP)^s843Tg^* individuals at 2, 3, and 5 dpf. N=3 independent experiments. ** p<0.01 in a two-sample t-test with Welch’s post hoc analysis for mean comparison. **D-E.** Expression level of *sting* in **D.** GFP+ ECs and **E.** GFP-cells (Other cells) sorted from Ctrl and EC-KD *Tg(kdrl:GFP)^s843Tg^* individuals at 2,3, and 5 dpf. N=3 independent experiments. * p<0.05; ** p<0.01 in a two-sample t-test with Welch’s post hoc analysis for mean comparison.

**Extended Figure 7.**
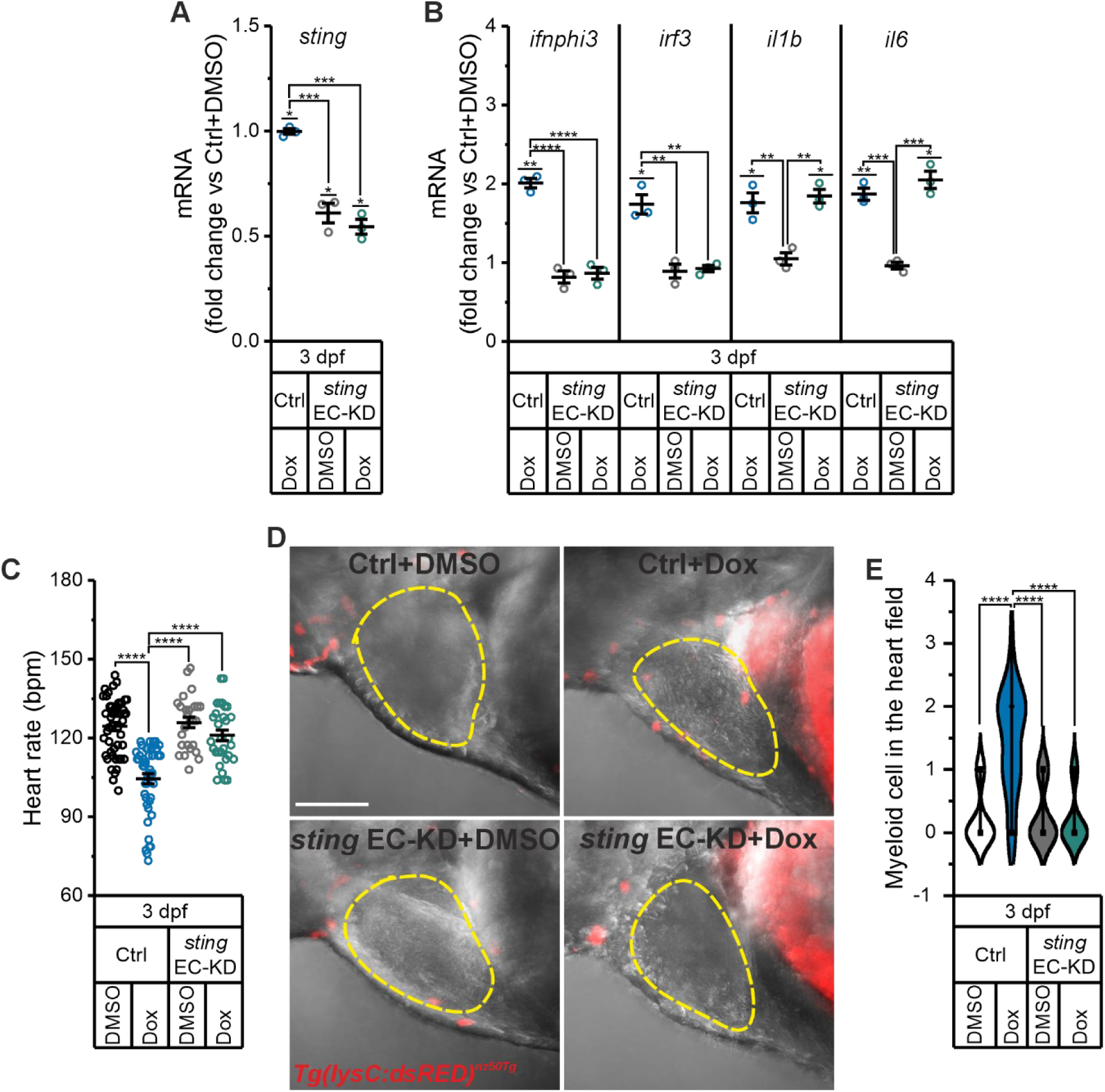
Loss of endothelial *rrm2b* confers protection from doxorubicin cardiotoxicity. **A-B.** Expression levels of **A.** *sting* and **B**. inflammatory genes determined by qRT-PCR in GFP+ ECs sorted from Ctrl and *sting* EC-KD *Tg(kdrl:GFP)^s843Tg^*individuals at 3 dpf with or without Dox treatment. N=3 independent experiments. * p<0.05; ** p<0.01; *** p<0.001; **** p<0.0001 in a one-way ANOVA test with Tukey’s post hoc analysis for mean comparison. **C.** Heart rate analysis at 3 dpf in Ctrl and *sting* EC-KD with or without Dox treatment. Bpm, beats per minute. Ctrl+DMSO (n=49), *sting* EC-KD+DMSO (n=26), Ctrl + Dox (n=46), *sting* EC-KD + Dox (n=30). N=3 independent experiments. **** p<0.0001 in a Kruskal-Wallis ANOVA test with Dunn’s post hoc analysis for mean comparison. **D.** Representative confocal image of the heart and myeloid cells in embryos from the transgenic line *Tg(lysC:dsRed2)^nz50Tg^* at 3 dpf. Red dots indicate the dsRED+ myeloid cells, while dotted lines contour the heart. Scale bar 100 µm. **H.** Quantification of myeloid cells in fish heart. Ctrl+DMSO (n=22), *sting* EC-KD+DMSO (n=21), Ctrl + Dox (n=22), *sting* EC-KD + Dox (n=20). N=3 independent experiments. **** p<0.0001 in a Kruskal-Wallis ANOVA test with Dunn’s post hoc analysis for mean comparison.

**Extended Figure 8.**
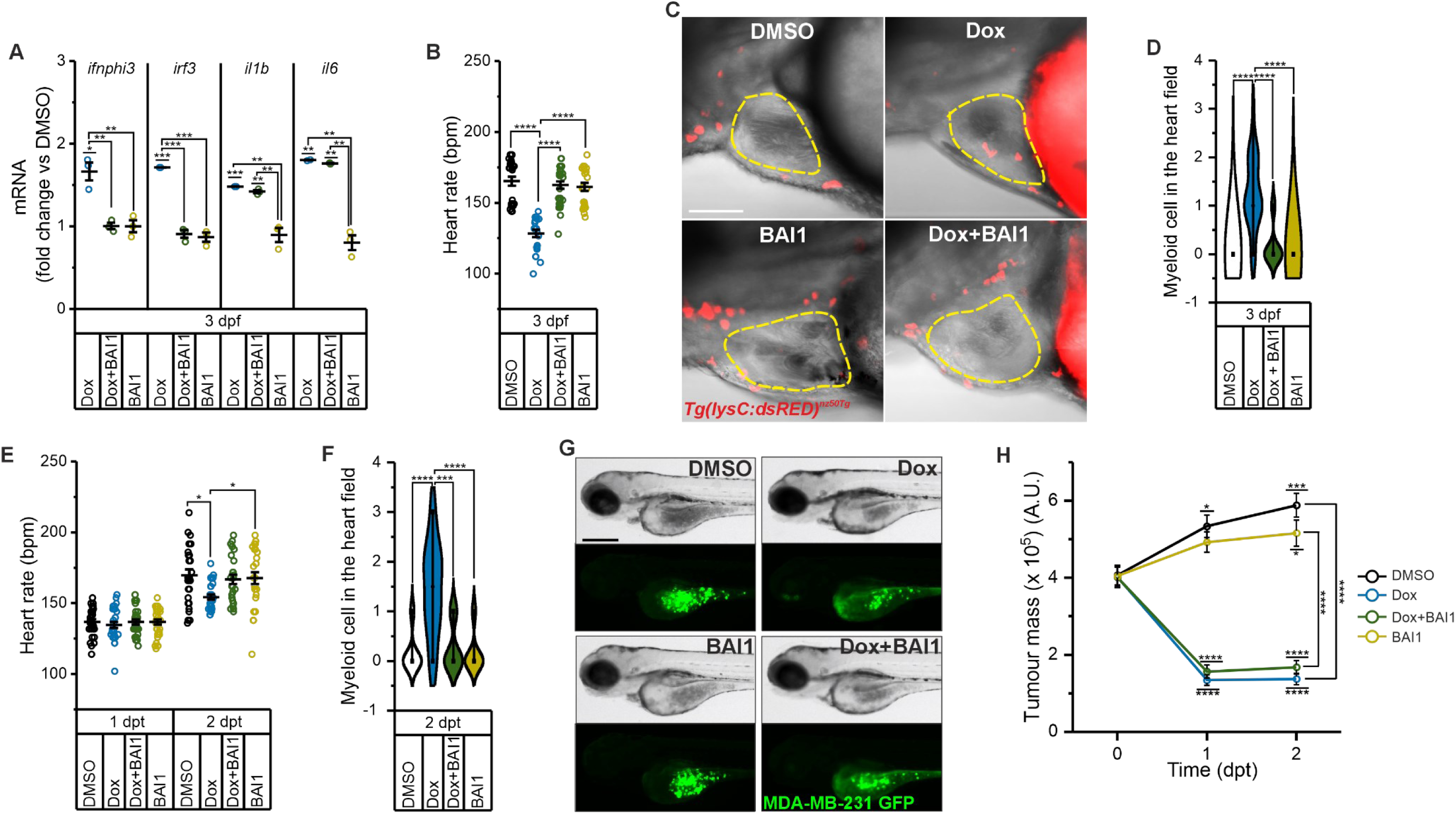
BAI1 prevents doxorubicin-induced cardiotoxicity while preserving its antitumor efficacy. **A.** Expression level of the indicated inflammatory genes in GFP+ sorted ECs from Dox-treated and DMSO control embryos, with or without the BAX inhibitor, BAI1, of the transgenic line *Tg(kdrl:GFP)^s843Tg^* at 3 dpf. N=3 independent experiments. * p<0.05; ** p<0.01; *** p<0.001 in a one-way ANOVA test with Tukey’s post hoc analysis for mean comparison. **B.** Heart rate analysis in embryos treated as indicated at 3dpf. Bpm, beats per minute. DMSO (n=24), Dox (n=23), Dox + BAI1 (n=28), BAI1 (n=22). N=3 independent experiments. **** p<0.0001 in a Kruskal-Wallis ANOVA test with Dunn’s post hoc analysis for mean comparison. **C-D.** Analysis of myeloid cell recruitment in the heart of embryos from the transgenic line *Tg(lysC:dsRed2)^nz50Tg^* treated as indicated at 3 dpf. **C.** Representative confocal image. Red dots indicate the dsRED+ myeloid cells, while dotted lines contour the heart. Scale bar 100 µm. **D.** Quantification of myeloid cells in fish heart. DMSO (n=21), Dox (n=23), Dox + BAI1 (n=23), BAI1 (n=21). N=3 independent experiments. **** p<0.0001 in a Kruskal-Wallis ANOVA test with Dunn’s post hoc analysis for mean comparison. **E-H.** Human breast cancer (MDA-MB-231) xenograft zebrafish model. Tumour cells were injected at 1 dpf and treated with Dox, or BAI or a combination of both drugs (0 dpt, days post treatment). Dox was removed after 24h (1 dpt) whereas BAI1 was maintained in fish water. Analyses were performed at 0, 1 and 2 dpt. **E.** Heart rate analysis. Bpm, beats per minute. DMSO (n=30), Dox (n=30), Dox + BAI1 (n=30), BAI1 (n=30) at 1dpt; DMSO (n=27), Dox (n=28), Dox + BAI1 (n=26), BAI1 (n=27) at 2dpt. N=3 independent experiments. * p<0.05 in a Kruskal-Wallis ANOVA test with Dunn’s post hoc analysis for mean comparison or in a one-way ANOVA test with Tukey’s post hoc analysis for mean comparison. **F.** Quantification of myeloid cells in fish heart of 2 dpt embryos from the transgenic line *Tg(lysC:dsRed2)^nz50Tg^* treated as described. DMSO (n=18), Dox (n=18), Dox + BAI1 (n=18), BAI1 (n=18). N=3 independent experiments. **** p<0.0001 in a Kruskal-Wallis ANOVA test with Dunn’s post hoc analysis for mean comparison. **G.** Representative brightfield and GFP images of the tumour mass in xenografted fish at 2 dpt. **H.** Time curve analysis of the tumour mass in xenografted fish. A.U, arbitrary units. DMSO (n=48), Dox (n=47), Dox + BAI1 (n=45), BAI1 (n=46) at 0 dpt, DMSO (n=42), Dox (n=36), Dox + BAI1 (n=37), BAI1 (n=44) at 1 dpt, DMSO (n=38), Dox (n=32), Dox + BAI1 (n=37), BAI1 (n=39) at 2 dpt. N=3 independent experiments. * p<0.05; ** p<0.01; *** p<0.001; **** p<0.0001 in a Kruskal-Wallis ANOVA test with Dunn’s post hoc analysis for mean comparison.

---

**Supplementary table 1.**

| REAGENTS |  |  |  |  |
| --- | --- | --- | --- | --- |
| ANTIBODIES |  |  |  |  |
|  | Source | Identifier | Host | Dilution |
| anti-HSP60 | Abcam | AB46798 | rabbit | 1:400 |
| anti-dsDNA | Abcam | AB27156 | mouse | 1:3000 |
| anti-BrdU (BrdU kit) | Roche | 11296736001 | mouse | 1:30 |
| anti-Lamin B1 | Santa Cruz Biotechnology | SC-6216 | goat | 1:1000 |
| anti-CYCLIN A | Sigma Aldrich | C4710 | mouse | 1:2000 |
| anti-p21 | SantaCruz Biotechnology | SC-817 | mouse | 1:500 |
| anti-Actin | Sigma Aldrich | A5316 | mouse | 1:4000 |
| anti-Vinculin | Sigma Aldrich | V9264 | mouse | 1:4000 |
| anti-TOM20 | Proteintech | 11802-1-AP | rabbit | 1:2000 |
| anti-HIS H3 | Cell Signaling technology | 9715S | rabbit | 1:4000 |
| anti-RRM2B | Abcam | ab-57217 | rabbit | 1:2000 |
| anti-RRM2 | Santa Cruz Biotechnology | SC-10844 | goat | 1:2000 |
| anti-p53 | Santa Cruz Biotechnology | SC-263 | mouse | 1:2000 |
| anti-TFAM | Cusabio | CSB-PA023413LA01Hu | rabbit | 1:1000 |
| HRP anti goat igG | Thermofisher | A15999 | donkey |  |
| HRP anti mouse igG | WesternSure | 926-80010 | goat |  |
| HRP anti rabbit igG | WesternSure | 926-80011 | goat |  |
| Alexa Fluor™ 568 anti-rabbit IgG | Thermofisher | A10042 | donkey | 1:500 |
| Alexa Fluor™ 488 anti-mouse | Thermofisher | A21202 | donkey | 1:500 |

| CELL CULTURE REAGENTS |  |  |
| --- | --- | --- |
|  | Source | Identifier |
| Endothelial Cell Growth Medium 2 Kit | Promocells | C-22111 |
| Endothelial Cell Growth Medium MV 2 kit | Promocells | C-22121 |
| DMEM, high glucose, GlutaMAX™ Supplement, pyruvate | Gibco | 10567022 |
| Fetal bovin serum | Gibco | A5256701 |
| Trypsin-EDTA 0.25%, phenol red | Gibco | 11560626 |
| DharmaFECT™4 | Dharmacon™ | T-2004-03 |
| Ethidium bromide | Sigma Aldrich | E1510 |
| Sodium pyruvate | Sigma Aldrich | P8574 |
| Uridine | Sigma Aldrich | U3003 |
| Penicillin-Streptomycin | Gibco | 11528876 |
| G-418 | Sigma-Aldrich | G8168 |

| CHEMICALS |  |  |
| --- | --- | --- |
|  | Source | Identifier |
| DMSO | Sigma-Aldrich | 41639 |
| Doxorubicin | Sigma-Aldrich | D1515 |
| Hydroxyurea | Sigma-Aldrich | H8627 |
| H151 | Invivogen | 941987-60-6 |
| BAI1 | MedChem Express | HY-103296 |
| Eltrobopag | MedChem Express | HY-15306 |
| EDTA | Sigma-Aldrich | E9884 |
| PBS | Gibco | 18912-014 |
| SDS | Sigma-Aldrich | 71729 |
| NaCl | Sigma-Aldrich | S7653 |
| HEPES | Gibco | 15630-122 |
| Digitonin | Sigma-Aldrich | D141 |
| Trizma base | Sigma-Aldrich | T1503 |
| Paraformaldehyde | Sigma-Aldrich | P6148 |
| RNAse A | Qiagen | 158922 |
| Proteinase K Solution | Invitrogen | 25530049 |
| Triton X-100 | Sigma-Aldrich | 93443 |
| Tween 20 | Sigma-Aldrich | P9416 |
| MAXBlock™ Blocking medium | Active Motif | 15252 |
| Prolong™ Gold antifade with DAPI | Invitrogen | P36931 |
| Glycin | Sigma-Aldrich | G8898 |
| Ethanol | VWR | 20821,31 |
| TRIzol Reagent | Thermo Fisher Scientific | 15596018 |
| HOT FirePol EvaGreen qPCR Supermix 5× | Solis BioDyne | 08-36-00001-10 |
| Taq Man™ Universal PCR Master Mix | Applied Biosystem | 4304437 |
| RIPA buffer | Sigma Aldrich | R0278 |
| phosphatase inhibitor | Roche | 4906845001 |
| protease inhibitor cocktail | Roche | 5892970001 |
| Bio-Rad Protein Assay Dye Reagent Concentrate | Biorad | 5000006 |
| 4-12% SDS-PAGE Gel | Invitrogen | NP0321 BOX |
| Immobilon® Forte or Immobilon® | Millipore | WBLUF500 |
| Classico Western HRP Substrates | Millipore | WBLUC0500 |
| Amersham™ Protran® nitrocellulose membrane for Western blot | Sigma | GE10600001 |
| Tricaine | Sigma-Aldrich | MS222 |
| 1-Phenyl 2-thiourea | Sigma-Aldrich | P7629 |
| Methyl cellulose | Sigma-Aldrich | M0387 |
| Agarose low melting | Sigma-Aldrich | A9414 |
| Collagenase type 2 | Wothington | LS004177 |
| Trypsin 0.5%, phenol red-free | Gibco | 15400054 |
| Opti-MEM | Gibco | 51985026 |

| KITS |  |  |
| --- | --- | --- |
|  | Source | Identifier |
| Senescent Detection Kit | Merck Millipore | KAA002 |
| BrdU Labeling and Detection kit I | Roche | 11296736001 |
| High-Capacity cDNA Reverse Transcription Kit | Applied Biosystem | 4374967 |
| Puregene Core kit A | Qiagen | 1042601 |
| NucleoSpin RNA XS kit | Macherey-Nagel | 740990.5 |
| Annexin V Apoptosis Detection Set PE-Cyanine7 | eBioscience | 25-8103-74 |
| TaqMan™ RNase P Control Reagents Kit | Applied Biosystems | 4316844 |

**Supplementary table 2.**
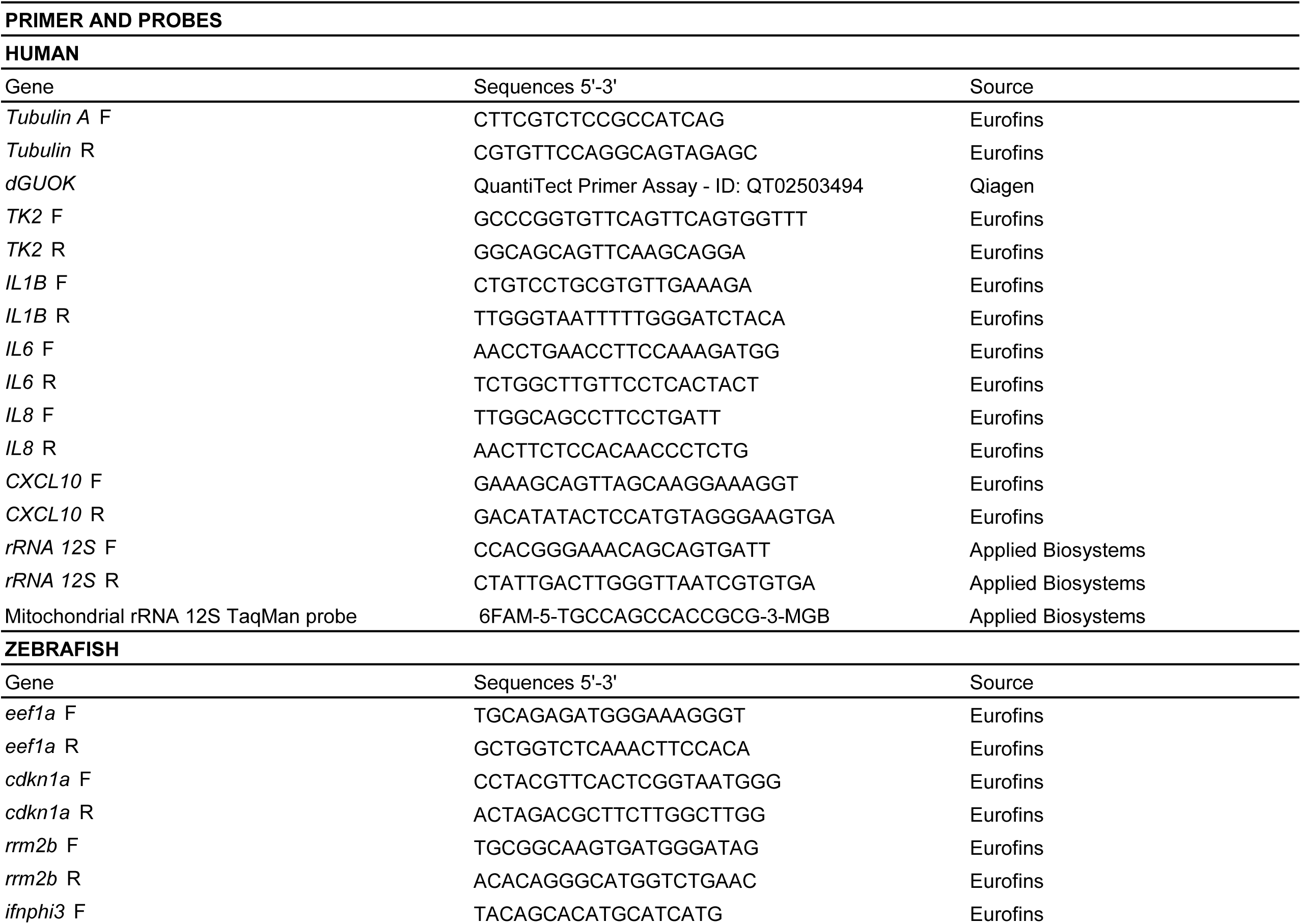
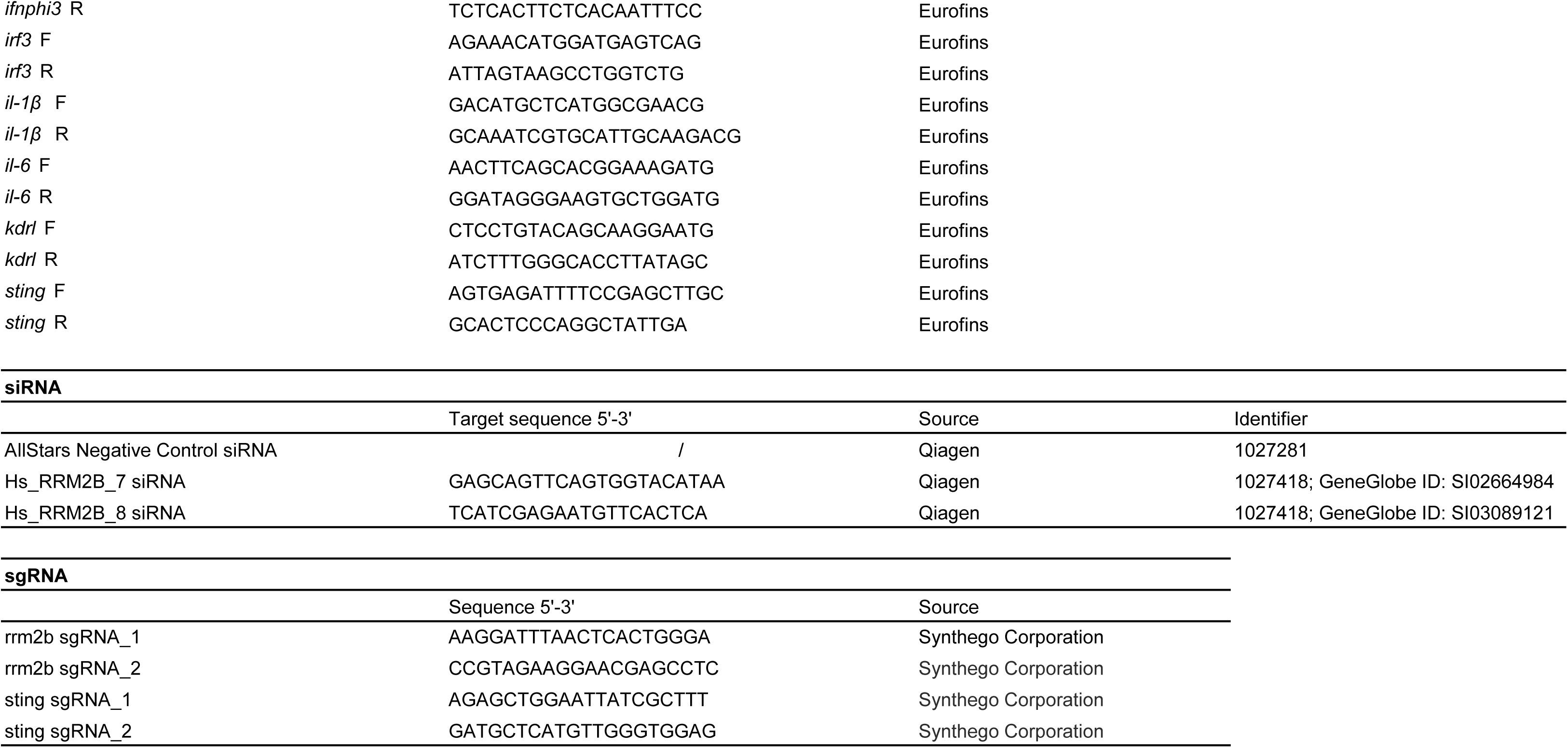

